# E-cadherin clustering as a regulator of morphogenesis

**DOI:** 10.64898/2026.04.20.719762

**Authors:** Gerald Lerchbaumer, Sergio Simoes, Ermia Etemadi, Fadi Zidan, Gonca Erdemci-Tandogan, Ulrich Tepass

## Abstract

Cell adhesion enables animal multicellular development. E-cadherin and the cadherin-catenin adhesion complex at adherens junctions are engaged in dynamic interactions with actomyosin generated contractile forces to drive epithelial morphogenesis. However, our understanding of how adhesion is regulated and how the tuning of adhesion contributes to morphogenesis remains incomplete. One key determinant of E-cadherin adhesion strength is clustering of the cadherin-catenin adhesion complex, a property studied extensively in vitro. Here, we use optogenetics to enhance E-cadherin cluster formation in the Drosophila embryo. Enlarged clusters were associated with increased E-cadherin surface abundance, assembled a normal cadherin-catenin complex, and showed reduced membrane mobility and turnover consistent with an increase in cell adhesion strength. Drosophila embryos with enhanced E-cadherin clustering displayed a severe reduction in cell intercalation and convergent extension of the anterior-posterior axis. To account for these observations, we modified existing vertex models to include junction-specific viscous forces representing E-cadherin-mediated friction between cells. This dissipative adhesion model predicts that enhanced adhesion increases resistance to cell rearrangements, thereby reducing cell neighbor exchanges and impairing convergent extension. To test model predictions, we analyzed two types of morphogenetic movements in embryos with enhanced E-cadherin clustering. Neuroblast ingression, which requires both apical constriction and cell rearrangement, was severely slowed. In contrast, mesoderm invagination, which requires apical constriction without neighbor exchanges, proceeded normally. Our findings suggest that optogenetic clustering, in contrast to overexpression of E-cadherin, is a valuable tool to examine the consequences of enhancing adhesion strength in tissue morphogenesis. Moreover, we propose that regulating E-cadherin clustering is essential for movements that require cell-cell contact changes.

## Introduction

During morphogenesis, epithelial sheets generate, transmit, and dissipate mechanical stress, but also maintain structural cohesion despite cell rearrangements. Tissues behave as viscoelastic materials in many instances, capable of flowing at short timescales but undergoing irreversible deformation under sustained stress [Lenne and Trivedi 2022; Noordstra et al., 2023; Campàs, et al., 2024; Boutillon et at., 2024]. Actomyosin-generated tension and E-cadherin-mediated cell adhesion at adherens junctions (AJs) are two main facilitators of these dynamic changes [Lecuit and Yap 2015; Yap et al., 2018; Agarwal and Zaidel-Bar 2021; Valet et al., 2022; Campàs et al., 2024]. Tension and adhesion are linked through feedback loops, as the cadherin-catenin complex (CCC) requires stabilization through cortical tension and contractility of the cell cortex and regulates dynamic remodeling of cell adhesion [Buckley et al. 2014; Lecuit and Yap, 2015; Cavanaugh et al., 2020; Sheppard et al., 2023; Campàs et al., 2024; Boutillon et at., 2024].

Both experimental and theoretical studies forecast an instructive role for cell adhesion in regulating tissue mechanics and guiding morphogenesis [Mongera et al. 2018; Iyer et al. 2019; Nestor-Bergmann et al. 2022; Sheppard et al., 2023]. The amount of cell adhesion is predicted to significantly influence the mechanical threshold, the ‘yield stress’, that tissues must overcome to undergo cell contact or shape changes [Nestor-Bergmann et al., 2022; Campàs et al., 2024; Boutillon et al., 2024]. Yield stress is defined by material properties that depend on the apico-lateral AJ, the zonula adherens, in particular in epithelial sheets that lack supportive ECM such as the Drosophila ectoderm. Theoretical models, especially vertex models of epithelial tissues, anticipate that modulating adhesion strength will alter tissue fluidity and the energy barriers for cell rearrangements in epithelia, though the relationship is complex [Lawson-Keister and Manning, 2021, Sahu et al., 2020]. In zebrafish embryos for example, tissue fluidization has been linked to a reduction in tissue yield stress, lowering the threshold for cell rearrangements and tissue flow [Mongera et al., 2018]. In Drosophila, evidence has highlighted the crucial role of dynamic and mechanoresponsive E-cadherin turnover for tuning tissue viscoelasticity and fluidization. Examples include imaginal disc morphogenesis [Iyer et al. 2019; Founounou et al. 2021], and cell intercalation that supports either tracheal tube elongation [Shaye et al., 2008; Shindo et al., 2008; Förster and Luschnig, 2012; Warrington et al., 2013] or body axis elongation in the Drosophila embryo [Rauzi et al., 2010; Levayer et al., 2011; Chandran et al., 2023]. However, how adhesion strength is controlled and how its modification impacts different morphogenetic movements remains an area of active research.

Actomyosin generated tension can be experimentally manipulated in vivo to either decrease or increase tension [Jordan and Karess, 1997; Winter et al., 2001; Simoes et al., 2010; Vasquez et al., 2014; Guglielmi et al., 2015; Izquierdo et al., 2018; Herrera-Perez et al., 2021]. In contrast, modification of CCC-mediated cell adhesion has largely been limited to loss-of-function approaches. It has not been possible to effectively increase E-cadherin adhesion strength and study its consequences in tissue morphogenesis. In vitro studies have identified E-cadherin clustering as an important determinant of E-cadherin function that enhances adhesion strength [Yap et al., 2015; Troyanovsky, 2023]. Here, we took advantage of an optogenetic method known as LARIAT [Lee et al., 2014] to induce cis-clustering of E-cadherin in vivo, causing a significant enhancement of adhesion strength, and examined its consequences on tissue morphogenesis. This optogenetic approach allowed us to further assess the role of adhesion regulation in tissue morphogenesis. Motivated by our experimental observations, we extended an anisotropic vertex model to include explicit viscous friction from cell adhesion. This dissipative adhesion model successfully reproduces our experimental phenotypes and suggests a mechanistic explanation for how adhesion dynamics regulates different morphogenetic movements.

## Results

### Optogenetic induction of E-cadherin clustering

Cadherins can form cis- and trans-clusters which are essential for effective adhesive interactions [Wu et al. 2010; Wu et al., 2015; Yap et al., 2015; Chen et al., 2015; Troyanovsky, 2023; Trojanovsky et al., 2025]. Clustering mechanisms were proposed to impact morphogenetic events in insect- and vertebrate axis elongation although evidence for a role in the regulation of morphogenesis remains indirect and limited [Levayer and Lecuit, 2013; Quang et al. 2013; Huebner et al. 2021]. We took advantage of the optogenetic LARIAT system [Lee et al., 2014] to promote clustering of E-cadherin (Fig. 1A-B). LARIAT is composed of Arabidopsis CRY2 fused to a nanobody targeting GFP (vhh) and CIBN fused to a multimerization domain [Lee et al., 2014]. LARIAT was activated by blue light in embryos expressing only endogenously GFP-tagged E-cadherin (endo-Ecad::GFP) in which LARIAT was maternally deposited (Ecad-LARIAT embryos) (Fig. 1C). As a control, we applied LARIAT to a GFP-tagged membrane-associated PH domain which formed large cytoplasmic clusters (Fig. 1C). In contrast, we observed a pronounced junctional accumulation of E-cadherin when clustering was induced globally in the embryo (Fig. 1C,D; Fig. S1A; Video S1). Junctional E-cadherin levels were increased by 2 to 3-fold compared to endo-Ecad::GFP controls (Fig. 1F). Quantitative assessment of images from structural illumination microscopy (SIM) revealed that 3D-cluster volume was significantly increased in Ecad-LARIAT embryos, with cluster distribution skewed towards larger less dense clusters (Fig. 1E,G,H). These findings suggest that LARIAT-mediated clustering drives the assembly of enlarged cis-clusters of E-cadherin within the plasma membrane. Interestingly, expressing the anti-GFP-CRY2 nanobody-fusion by itself (Ecad-CRY2 embryos) caused a degree of E-cadherin clustering that was intermediate between endo-Ecad::GFP control embryos and Ecad-LARIAT embryos (Fig. 1A,C,F), resulting from the light-dependent homo-oligomerization of CRY2 [Bugaj et al., 2013; Lee et al., 2014]. This offers an elegant, dose dependent approach to tune E-cadherin clustering in living tissues. Our observations suggest that E-cadherin can be effective clustered in the plane of the membrane, potentially increasing its activity in adhesion rather than disrupting its function.

**Figure 1:**
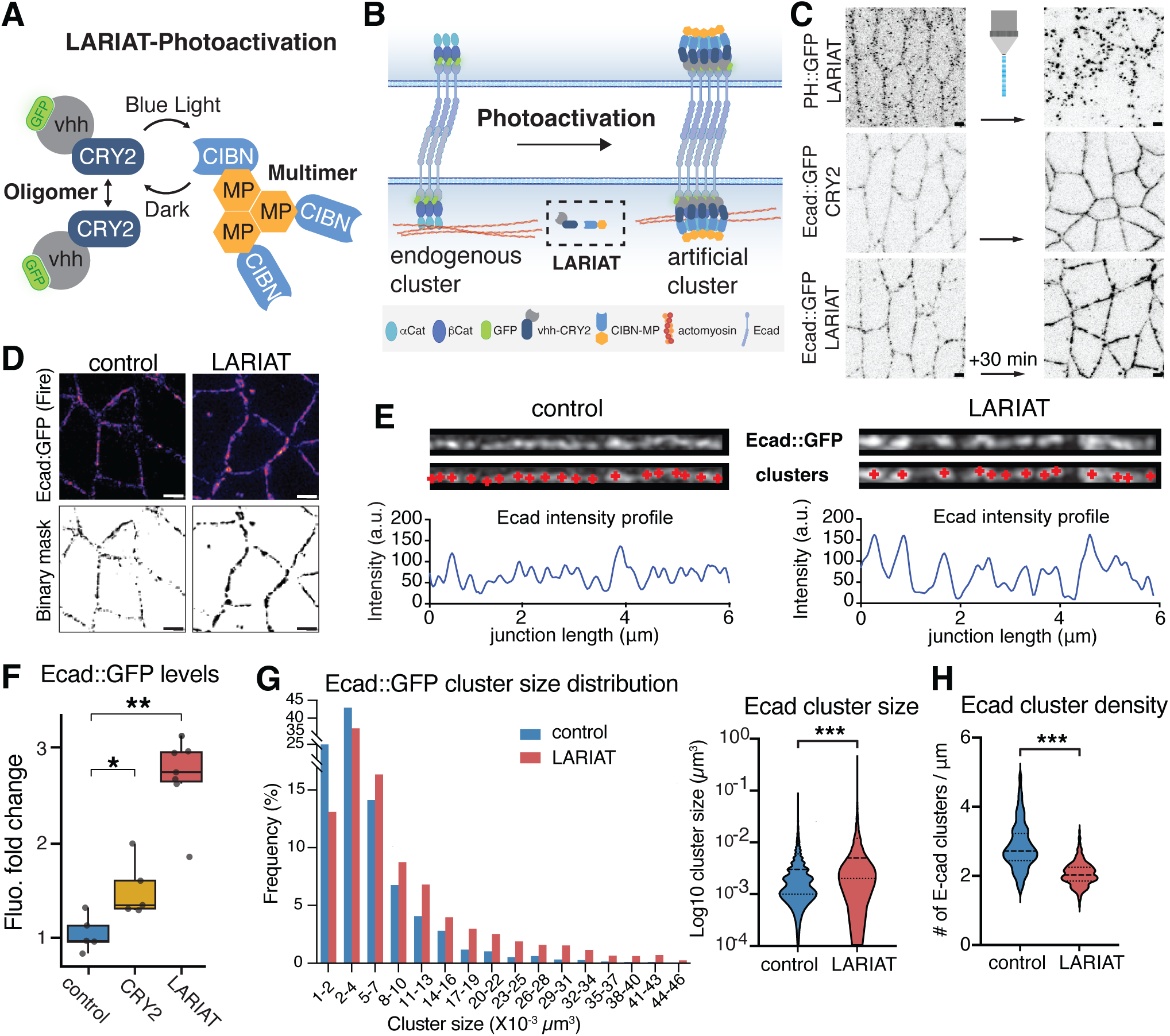
Light-induced clustering increases cluster size and membrane levels of E-cadherin. **(A)** Schematic illustration of the LARIAT system. A GFP-binding nanobody (vhh) is fused to CRY2; CIBN is fused to a multimerization domain (MP). Blue-light exposure causes CRY2 to interact with CRY2 and CIBN. MP-multimers form constitutively. **(B)** Schematic illustration of LARIAT with endogenously tagged E-cadherin (endo-Ecad::GFP). Blue-light exposure drives the formation of enlarged CCC clusters in the plasma membrane. **(C)** Confocal images showing LARIAT applied to PH::GFP, which clusters predominantly in the cytoplasm after 30 minutes of blue light-activation. Ecad::GFP clusters at adherens junctions with CRY2 only or LARIAT. Scale bars, 2 µm. **(D)** MI-SIM images of Ecad::GFP in control and Ecad-LARIAT embryos (st. 8, germband). Fire LUT images and corresponding binary masks after thresholding. Scale bars, 2µm. **(E)** Ecad::GFP fluorescence by MI-SIM of a single straightened bicellular junction in control and Ecad-LARIAT embryos (st. 8, germband). Red plus signs indicate the position of local fluorescence maxima (individual E-cadherin clusters), illustrated by a fluorescence intensity profile below and quantified in (H). **(F)** Quantification of perimeter fluorescence intensity shows an increase in junctional Ecad::GFP levels in Ecad-CRY2 and Ecad-LARIAT embryos. Wilcoxon rank-sum test: control vs vhhGFP, p = 0.0317; control vs LARIAT, p = 0.00253. Sample sizes: control = 5 embryos, CRY2 only = 5 embryos, LARIAT = 7 embryos. **(G)** Binned Ecad::GFP cluster size distribution in control and Ecad-LARIAT embryos (st. 8, germband); Ecad::GFP cluster sizes are significantly increased in Ecad-LARIAT embryos. Median/Q1/Q3 cluster volumes (µm³): 0.003/0.001/0.006 in controls versus 0.005/0.002/0.012 in LARIAT embryos (n = 5984 clusters in 15 control embryos; n = 3567 clusters in 11 LARIAT embryos). Wilcoxon rank-sum test: control vs LARIAT p = 6.01 × 10⁻⁷⁴. **(H)** E-cadherin cluster density (# of clusters per μm of junctional length) in control and LARIAT embryos. Each data point indicates one individual junction. n = 174 junctions in 11 embryos (control) and n = 230 junctions in 11 embryos (LARIAT).

An alternative strategy to increase E-cadherin surface abundance and potentially cell adhesion is through overexpression. We therefore compared LARIAT-induced E-cadherin clustering with the impact of E-cadherin overexpression. To maximize overexpression, we combined a ubiquitin promoter driven transgene (ubi-Ecad::GFP) with a UAS-driven transgene expressed by a strong maternal Gal4 driver (UAS-Ecad::GFP) in a wildtype background. These Ecad-OE embryos showed an increase in junctional and cytoplasmic Ecad::GFP levels at the onset of germband extension compared to endo-Ecad::GFP controls (Fig. S2A,C,D), as previously reported (Wang et al., 2024). However, Ecad-OE levels declined between 15 and 25 minutes after the onset of germband extension (Fig. S2D). In contrast, Ecad-LARIAT embryos controlled by the same maternal Gal4 driver displayed much higher E-cadherin levels (∼3 fold) that persistently increased over the observation period (Fig. S2D). These data suggest that Ecad-OE embryos robustly regulate E-cadherin membrane levels through protein trafficking whereas Ecad-LARIAT embryos have a reduced turnover rate (Fig. S2B). The efficient removal of E-cadherin from the membrane in Ecad-OE embryos is consistent with previous findings that E-cadherin overexpression has little effect on normal development [e.g. Pacquelet and Rorth, 2005; Wang et al., 2024], suggesting that overexpression alone does not generate a substantial adhesion gain-of-function phenotype. Accordingly, Ecad-OE embryos showed no defect in axis elongation (Fig. S2E). To quantify tissue extension, we compared cell displacement between control, Ecad-OE, and Ecad-LARIAT embryos (Fig. S2E). Control and OE embryos displayed similar tissue extension, whereas Ecad-LARIAT embryos showed a significant reduction (see below for further discussion on the effect of E-cadherin clustering on germband extension). Together, these data suggest that the robust increase in junctional E-cadherin levels and altered junctional dynamics in Ecad-LARIAT embryos result in a strong increase in adhesion strength that cannot be reproduced by E-cadherin overexpression.

### Clustering of E-cadherin reduces protein mobility and turnover and enhances cell adhesion

We hypothesized that the enhanced protein levels and increased clustering of E-cadherin at AJs in Ecad-LARIAT and Ecad-CRY2 embryos causes an effective increase in cell adhesion. E-cadherin mobility and turnover are thought to be important mechanisms of orienting contractility and tuning cell adhesion during tissue rearrangements [Levayer and Lecuit 2013; Malinverno et al. 2017; Iyer et al. 2019; Chandran et al. 2021; Founounou et al. 2021]. Turnover-rates of adhesion receptors such as E-cadherin have been correlated with tissue plasticity and viscoelasticity in developmental contexts, whereby morphogenetically stable tissues show comparatively low turnover [Huang et al., 2011, Iyer et al., 2019, Zhang et al., 2024]. Conclusions about a causal relationship between tissue properties and E-cadherin turnover remain limited, as most studies report correlations rather than direct perturbations. Moreover, experimental manipulation of E-cadherin turnover has typically relied on indirect approaches, such as global inhibition of endocytosis, which can produce pleiotropic effects [e.g. Cavey et al. 2008; Levayer et al., 2010; Tamada et al., 2012].

To assess cell-cell adhesion in Ecad-LARIAT embryos we characterized the dynamic properties of E-cadherin. We examined E-cadherin lateral mobility and turnover using fluorescent recovery after photobleaching (FRAP) analysis in Ecad-LARIAT embryos during gastrulation when junctional E-cadherin levels are regulated in support of cell intercalation [Blankenship et al., 2006; Simoes et al., 2010; Levayer et al., 2011; Levayer and Lecuit 2013] and cell ingression [Simoes et al., 2017]. We carried out two distinct photobleaching experiments: in one set, we bleached a small spot (∼ 1 *μ*m) within a bicellular junction (BCJ) (spot FRAP), and in a second set of experiments we bleached the entire BCJ between two tricellular junctions (TCJs; whole junction FRAP) (Fig. 2A).

**Figure 2.**
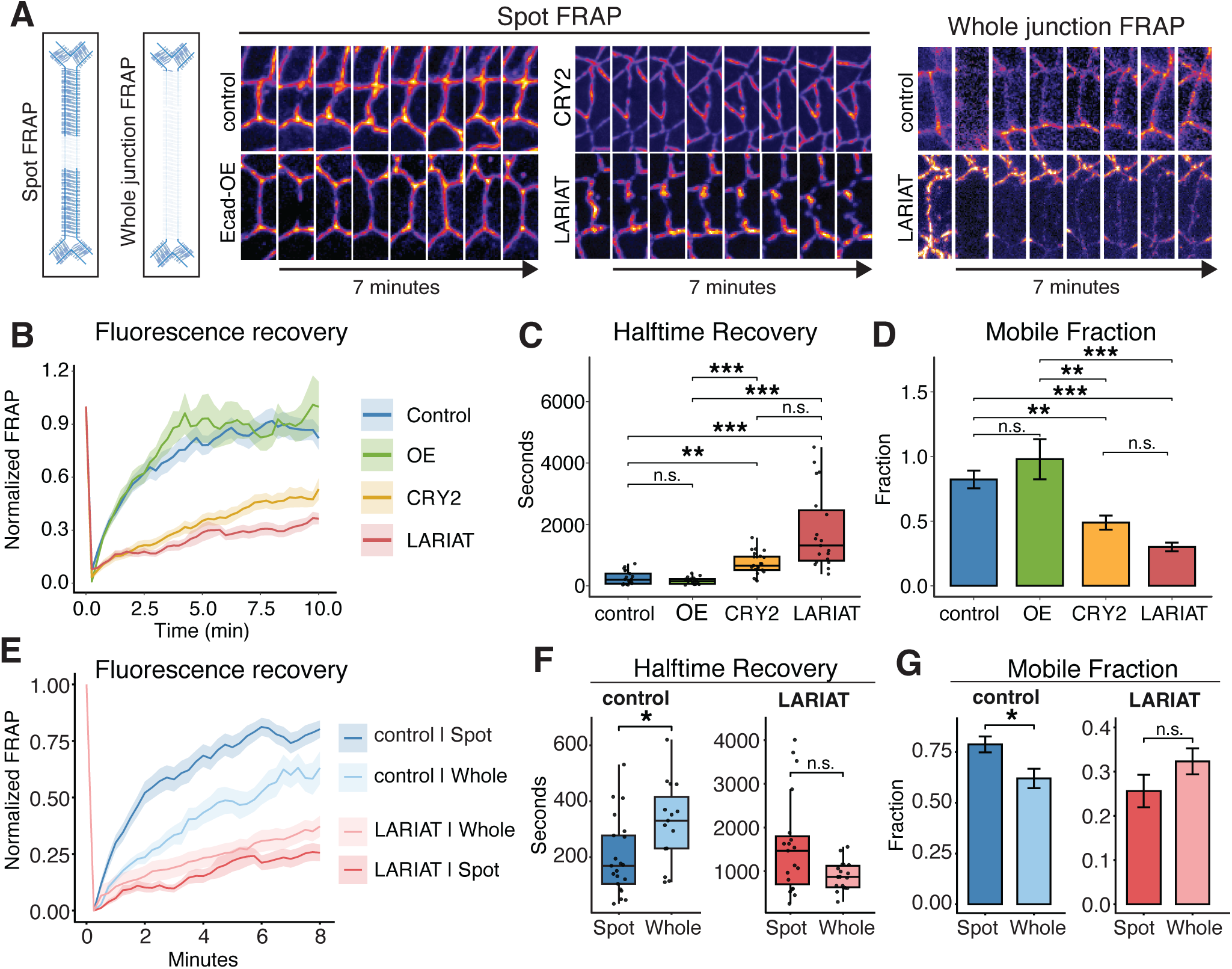
Optogenetic clustering reduces E-cadherin recovery and lateral mobility. **(A)** Schematic illustration of the spot FRAP and whole-junction FRAP assays, and representative spot FRAP and whole-junction FRAP kymograph. **(B-D)** Fluorescence recovery curves from spot FRAP for control, Ecad overexpression (Ecad-OE), Ecad-CRY2, and Ecad-LARIAT conditions, with corresponding quantification of recovery half-time (C) and mobile fraction (D). Recovery was strongly reduced in Ecad-LARIAT embryos and was also altered in Ecad-CRY2 embryos relative to control, whereas Ecad-OE remained similar to control. Mean recovery half-times were 252.0 ± 47.2 s in control, 160.4 ± 27.4 s in Ecad-OE, 734.4 ± 74.5 s in Ecad-CRY2, and 1793.7 ± 296.7 s in Ecad-LARIAT embryos. Mean mobile fractions were 0.822 ± 0.069 in control, 0.979 ± 0.155 in Ecad-OE, 0.489 ± 0.054 in Ecad-CRY2, and 0.301 ± 0.034 in Ecad-LARIAT embryos. Sample sizes were: control 21 junctions, Ecad-OE 17 junctions, Ecad-CRY2 23 junctions, LARIAT 19 junctions. Statistics: Kruskal-Wallis test with Dunn’s multiple-comparisons test and Bonferroni correction. For recovery halftime: control vs Ecad-OE, n.s. (p = 1.0); Control vs Ecad-CRY2, p = 9.66 × 10^-4; control vs LARIAT, p = 9.45 × 10^-8; Ecad-OE vs Ecad-CRY2, p = 1.14 × 10^-4; Ecad-OE vs LARIAT, p = 8.85 × 10^-9; Ecad-CRY2 vs LARIAT, n.s. (p = 0.214). For mobile fraction: control vs Ecad-OE, n.s. (p = 1.0); Control vs Ecad-CRY2, p = 5.52 × 10^-3; Control vs LARIAT, p = 7.50 × 10^-7; Ecad-OE vs Ecad-CRY2, p = 2.85 × 10^-3; Ecad-OE vs LARIAT, p = 4.33 × 10^-7; Ecad-CRY2 vs LARIAT, n.s. (p = 0.169). **(E-G)** Comparison of spot FRAP and whole-junction FRAP in control and Ecad-LARIAT embryos. Spot FRAP values are identical to those shown in (B). In control embryos, fluorescence recovered faster and to a higher plateau in spot FRAP than in whole-junction FRAP, consistent with a shorter recovery half-time (F) and higher mobile fraction (G). In Ecad-LARIAT embryos, recovery was strongly reduced in both assays. Mean recovery half-times were 252.0 ± 47.2 s and 323.6 ± 37.6 s for control spot and whole-junction FRAP, respectively, and 2024.6 ± 364.1 s and 894.3 ± 77.5 s for LARIAT spot and whole-junction FRAP. Mean mobile fractions were 0.822 ± 0.069 and 0.620 ± 0.048 in control, and 0.288 ± 0.034 and 0.323 ± 0.029 in LARIAT, for spot and whole-junction FRAP, respectively. In control embryos, spot and whole-junction FRAP differed significantly for both recovery half-time (Welch’s t-test, p = 0.0158) and mobile fraction (Welch’s t-test, p = 0.0105). In contrast, no significant differences were detected between spot and whole-junction FRAP in Ecad-LARIAT embryos (half-time, Wilcoxon rank-sum test, p = 0.0861; mobile fraction, Welch’s t-test, p = 0.1630).

Spot FRAP on endo-Ecad::GFP control embryos showed the expected dynamics, with rapid and exponential initial recovery over the first 2 minutes after bleach. Subsequent recovery slowed and plateaued after 5 minutes consistent with previous findings (Fig. 2B) [Bulgakova et al., 2013, Moreno et al., 2022]. This recovery curve is the result of two components: (i) lateral diffusion within the plasma membrane that causes the initial exponential increase in fluorescence and (ii) the turnover of E-cadherin through endocytosis and exocytosis [de Beco et al., 2009; Bulgakova et al., 2013; Bulgakova and Brown, 2016, Iyer et al., 2019]. Ecad-LARIAT and Ecad-CRY2 embryos showed a strikingly different recovery curve with a strongly reduced early exponential phase, and Ecad::GFP plateauing at significantly lower levels with a recovery half-time approximately 7.1times greater than control and a 63% reduced mobile fraction in Ecad-LARIAT embryos and 2.9 times greater than control and a 41% reduced mobile fraction in Ecad-CRY2 embryos (Fig. 2B-D). In contrast, spot FRAP on embryos overexpressing Ecad::GFP did not show a significant difference to fluorescent recovery compared to controls (Fig. 2B-D). Thus, membrane mobility and turnover were strongly reduced upon LARIAT induced E-cadherin clustering whereas Ecad-CRY2 embryos showed E-cadherin kinetics intermediate between Ecad-LARIAT embryos and controls.

Whole junction FRAP on control embryos showed a strong reduction in the rapid phase of fluorescent recovery compared to spot FRAP data, likely because E-cadherin does not readily diffuse across tricellular junctions (TCJs) (Fig. 2E). Consequently, we observed an increase in the half-time recovery in endo-Ecad::GFP controls compared to spot FRAP (Fig. 2F). The observed plateauing suggests a mobile fraction of ∼60% (Fig. 2G). In contrast, whole junction FRAP in Ecad-LARIAT embryos showed comparable recovery kinetics to spot FRAP results (Fig. 2E-G). To further assess lateral mobility of clusters in Ecad-LARIAT embryos we examined cluster movements in relation to a nearby TCJ over a 10-minute period. We found that cluster displacement is considerably reduced in Ecad-LARIAT embryos compared to controls (Fig. S3; Video S2) consistent with our FRAP data, further indicating that LARIAT-based clustering interferes with E-cadherin lateral mobility. These results demonstrate that LARIAT-mediated E-cadherin clustering strongly reduces lateral mobility of E-cadherin and E-cadherin clusters, and the turnover of E-cadherin.

### Ecad-LARIAT clusters recruit other CCC components

To further test whether Ecad-LARIAT embryos assemble functional AJs, we analyzed the recruitment of E-cadherin binding partners: α-Catenin (αCat), Armadillo (Arm, Drosophila β-catenin), Vinculin and F-actin (Fig. 3A). Live imaging of Ecad-LARIAT embryos expressing endo-αCat::RFP revealed a 2 to 3-fold increase of junctional E-cadherin levels accompanied by a ∼1.4-fold increase in αCat (Fig. 3A,B). Antibody staining for Arm in fixed Ecad-LARIAT embryos showed a fluorescence increase of 1.4-fold compared to controls (Fig. 3A,B). This result is corroborated by findings that expression of the LARIAT construct in Engrailed (En) stripes and global blue light activation is sufficient to induce Arm enrichment in the En domain (Fig. S4A). Interfaces between En positive and En negative cells showed an intermediate degree of enhanced E-cadherin clustering (Fig. S4B). αCat and Arm colocalize with large E-cadherin clusters (Figs. 3C, S4C), which was quantified by calculating the Spearman colocalization coefficient between E-cadherin and both catenins (Fig. 3D). We further confirmed significant colocalization of E-cadherin with Vinculin and F-actin in both control and LARIAT embryos (Figs. 3D, S4C).

**Figure 3:**
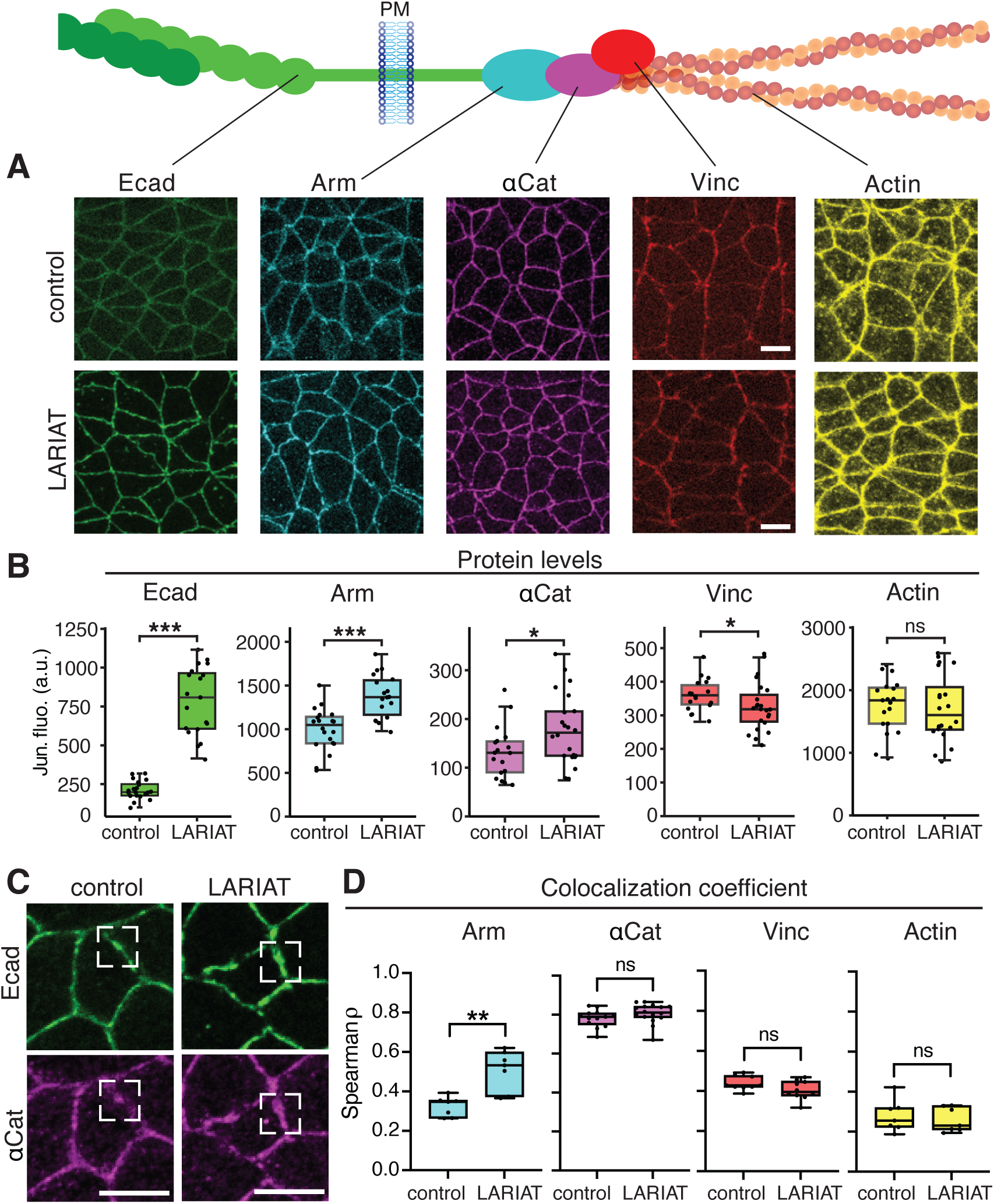
Cadherin-catenin complex is recruited into LARIAT induced E-cadherin clusters. **(A)** Illustration of the cadherin-catenin-complex bound to actin and images of respective proteins in control and Ecad-LARIAT stage 8 embryos. Scale bars, 5µm. **(B)** Quantification of membrane protein levels of CCC proteins reveals increased levels in Arm (= ß-Catenin) and αCat, reduced levels of Vinculin (Vinc) and no changes in actin in Ecad-LARIAT embryos. Dots represent individual embryos. Wilcoxon rank-sum test, Ecad p = 5.80 × 10^-11, Arm p = 5.18 × 10^-5, actin p = 0.749, αCat p = 0.0307, Vinc p = 1.16 × 10^-7. Sample sizes: Ecad, control = 20 embryos, LARIAT = 19 embryos; Arm, control = 20 embryos, LARIAT = 18 embryos; actin, control = 19 embryos, LARIAT = 20 embryos; αCat, control = 18 embryos, LARIAT = 22 embryos; Vinc, control = 18 embryos, LARIAT = 23 embryos. **(C)** Colocalization of E-cadherin and αCat at individual clusters in a control embryo and a Ecad-LARIAT embryo. Scale bar, 5µm. **(D)** Spearman colocalization coefficients between E-cadherin and other CCC proteins. Positive correlation was found between all tested pairs, with no significant differences between control and Ecad-LARIAT embryos, except for Arm with stronger colocalization in Ecad-LARIAT embryos. Box plots show median, interquartile range and min–max values. Each data point represents one junctional mask consisting of 29-65 junctions,1-3 masks per embryo. αCat: control: 598 junctions, 5 embryos; LARIAT: 682 junctions, 6 embryos. Arm and F-actin: control: 264 junctions, 7 embryos; LARIAT: 266 junctions, 7 embryos. Vinc: control: 520 junctions, 9 embryos; LARIAT: 543 junctions, 7 embryos. Negative controls (not shown) yielded near zero Spearman coefficients (median in control/LARIAT: -0.016/-0.008; max in control/LARIAT: 0.034/0.065; min in control/LARIAT: -0.067/-0.067 and inter-quartile range in control/LARIAT: -0.028: -0.0123/-0.019:0.0136). Statistics: two-tailed Mann-Whitney U test. ** p = 0.0023.

Together, our results indicate that optogenetically induced E-cadherin clusters are associated with E-cadherin binding partners, supporting the formation of large cadherin-catenin assemblies. Interestingly, we noticed a reduction of Vinculin recruitment to E-cadherin clusters in Ecad-LARIAT embryos (Fig. 3B). Vinculin recruitment and binding to αCat is tension-sensitive and depends on the local strength of actomyosin forces in the Drosophila embryo [Kale et al., 2018; Sheppard et al., 2023]. This observation raises the possibility that actomyosin generated forces in Ecad-LARIAT embryos are similar to controls but distributed over an increased number of E-cadherin adhesion complexes at the plasma membrane, leading to less force per CCC and a reduction in Vinculin recruitment.

Taken together, our findings support the conclusion that E-cadherin clustering is a key determinant of adhesion strength in vivo. Optogenetically induced E-cadherin clusters are competent to recruit binding partners and effectively engage actomyosin. Moreover, we showed that optogenetic clustering of E-cadherin reduces its lateral mobility and turnover significantly, increasing CCC abundance at AJs. By stabilizing E-cadherin at AJs in functional adhesion complexes, the LARIAT system provides a tool to increase adhesion strength and assess its impact on cell and tissue-scale mechanics during morphogenesis.

### Optogenetic clustering of E-cadherin has little impact on myosin activity and junctional tension

The CCC anchors actin to AJs and recruits modulators of myosin II activity [Desai et al., 2013; Buckley et al., 2014; Yashiro et al. 2014; Lecuit and Yap, 2015; Curran et al. 2017; Ishiyama et al., 2018; Sumi et al. 2018]. The role of myosin in junctional remodeling during epithelial morphogenesis has been investigated in the context of Drosophila germband extension [Zallen and Wieschaus 2004; Bertet et al., 2004; Fernandez-Gonzalez et al., 2009; Rauzi et al., 2010; Levayer and Lecuit 2013; Tetley et al., 2016], and in the Drosophila pupal notum [Curran et al., 2017; Sumi et al., 2018]. Myosin is enriched at shrinking (vertical) junctions during cell intercalations that drive germband extension, while E-cadherin is slightly enriched at stable horizontal junctions. Anisotropic shrinking of junctions is supported by flows of medial myosin towards the shrinking junctions. These flows appear to coincide with imbalances in E-cadherin endocytosis [Rauzi et al., 2010; Levayer et al., 2011; Levayer and Lecuit 2013]. Interestingly, our analysis revealed that optogenetic clustering of E-cadherin results in a shift of E-cadherin distribution from planar polarized to uniform, while levels and planar polarity of myosin were not affected (Fig. 4A-E, S5A). The persistence of myosin anisotropy suggests that AP planar patterning cues such as the Rho and Toll pathways [Levayer et al., 2011; Pare et al., 2014; Tamada et al., 2021] remain intact upon E-cadherin clustering and do not respond to differences between E-cadherin levels at vertical and horizontal junctions.

**Figure 4:**
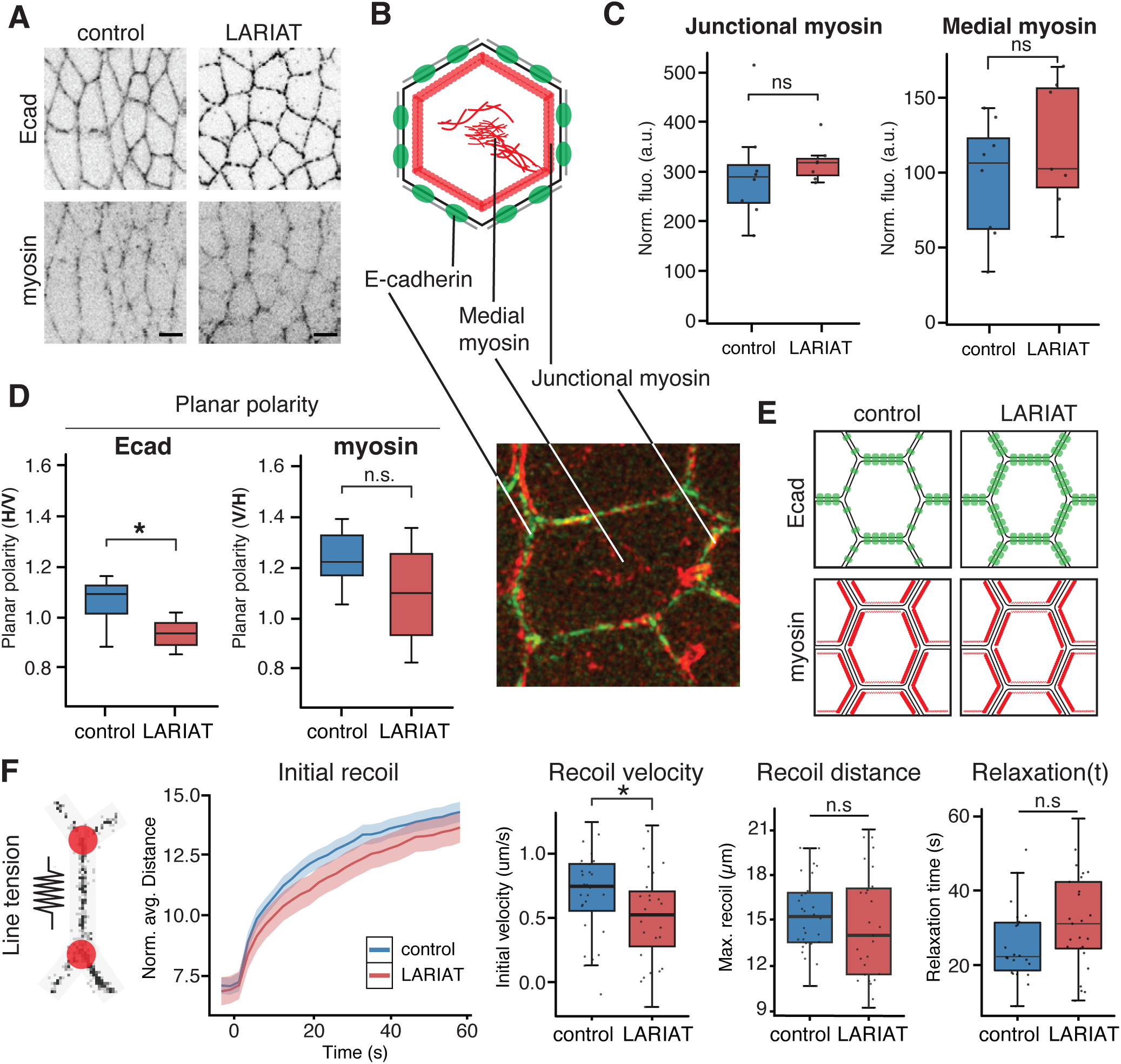
Optogenetic clustering of E-cadherin has little effect on myosin. **(A)** Images of endo-Ecad::GFP and sqh-Sqh::mCherry in the germband (stage 8) of control and Ecad-LARIAT embryos. Scale bars, 5 µm. **(B)** Illustration of E-cadherin, medial and junctional myosin in the apical domain. The same proteins are shown in SIM-image of an ectodermal cell at stage 9. **(C)** Quantification of junctional and medial myosin II levels in control and Ecad-LARIAT embryos. Junctional and medial myosin levels are unaffected by enhanced E-cadherin clustering. Dots are individual embryos. Statistics: Wilcox rank sum test, Jun. myosin. p = 0.3253, Med. myosin. p = 0.4519, Sample sizes: control = 8 embryos (337 cells), LARIAT = 7 embryos (308 cells). **(D)** Planar polarity of E-cadherin and myosin assessed as intensity ratios. The fluorescence ratio of horizontal-to-vertical (H/V) cell contacts of E-cadherin was significantly lowered in the Ecad-LARIAT embryos compared to control. Myosin planar polarity shown as vertical-to-horizontal (V/H) ratio was normal in Ecad-LARIAT embryos. Statistics: Wilcox rank sum test, Ecad p = 0.024, Myosin p= 0.183, Sample sizes: control = 8 embryos, 485 junctions, LARIAT = 7 embryos, 479 junctions. **(E)** Illustration of planar polarized distribution of E-cadherin and myosin during germband extension in control and Ecad-LARIAT embryos. **(F)** Recoil of tricellular junctions after laser ablation of vertical junctions shows a small reduction in junctional tension in Ecad-LARIAT embryos. Recoil distance within the first 60 seconds post-ablation is shown in a line-graph with shaded SEM ribbons. Initial recoil velocity measured immediately after ablation, showed a significant reduction in Ecad-LARIAT embryos. Measurements of recoil distance and tissue relaxation did not show significant differences between conditions. Statistics: Wilcoxon rank sum test, recoil velocity p = 0.049, recoil distance p = 0.4, relaxation time p = 0.068. Sample sizes: control: 12 embryos, 29 junctions, Ecad-LARIAT 13 embryos, 30 junctions.

Edge contractility and tension can be inferred from relative levels of myosin [Fernandez-Gonzalez et al., 2009; Curran et al. 2017; Streichan et al. 2018]. Surprisingly, comparison between control and Ecad-LARIAT embryos during germband extension did not reveal significant changes in myosin levels (Fig. 4A,C). Moreover, FRAP analysis of myosin recovery did not show significant changes in myosin turnover (Fig. S5B). Because of the potential contributions of medial myosin activity to junctional rearrangements [Rauzi et al., 2010; Levayer and Lecuit, 2013] we quantified myosin flows using particle image velocimetry (PIV). We found that Ecad-LARIAT embryos showed faster medial myosin flows and less planar polarized flows, i.e. medial myosin behavior appeared more isotropic (Fig. S5C). To further examine how E-cadherin clustering affects myosin activity, we performed laser ablation experiments on vertical junctions in the germband (Fig. 4F; Video S3). Comparing the initial recoil velocities of TCJs revealed that clustering of E-cadherin led to a marginal decrease in line tension (Fig. 4F). However, the maximum displacement of TCJs (recoil distance) and the tissue relaxation (viscosity) were not significantly altered (Fig. 4F). A lack of enhanced actomyosin induced tension is also supported by the observed reduction rather than increase in Vinculin recruitment to AJs, as mentioned [Kale et al., 2018; Sheppard et al., 2023] (Fig. 3B). Considering that junctional and medial myosin levels appeared normal in Ecad-LARIAT embryos despite the strong increase of the CCC at AJs, we hypothesize that the altered ratio between adhesion molecules and actomyosin is responsible for a decrease in tension force applied to AJs. Taken together, these findings suggest that clustering of E-cadherin and the associated increase in adhesion strength has a minor if any impact on myosin flows and little impact on levels and planar distribution of myosin.

### Stabilizing E-cadherin through clustering inhibits cell rearrangements

Myosin is the major force generator driving cell intercalations during germband extension [Zallen and Wieschaus 2004; Bertet et al., 2004; Blankenship et al. 2006]. One consequence of myosin contractility in the germband is that when AJs are compromised, cells detach from each other in a pattern that mimics the enrichment of myosin [Sawyer et al., 2010; Sheppard et al., 2023]. Given that myosin levels and dynamics appeared near-normal in Ecad-LARIAT embryos and only a small reduction in line tension was observed, we wondered whether the strong increase in CCC, its enhanced clustering, reduced turnover, and the associated increase in adhesion strength would impact cell rearrangements during tissue extension.

Notably, we found that germband extension was slow in Ecad-LARIAT embryos and the germband failed to fully extend compared to wild-type controls (Fig. 5A; Video S4 and S5). Posterior midgut invagination, which contributes to germband extension [Collinet and Lecuit, 2021] appeared normal. In contrast, the cell intercalations, known as T1 transitions [Kong et al., 2016; Pare and Zallen, 2020; Collinet and Lecuit, 2021] that are the major driving force of the convergent extension of the germband were dramatically reduced in Ecad-LARIAT embryos (Fig. 5B,C; Video S6). Overall, the number of T1 transitions per minute was reduced by ∼70% in Ecad-LARIAT embryos compared to controls (Fig. 5C), i.e. the average number of successful T1s was reduced from 4 T1 events per minute in control regions of interest to approximately 1 T1 per minute in Ecad-LARIAT embryos. Moreover, the remaining T1 transitions displayed slowed edge contraction of vertical junctions, junctions remained as 4-cell vertices for an increased amount of time, and horizontal junction expansion was delayed (Fig. 5D). Interestingly, in cases where new junctions formed in the LARIAT condition, we observed very low levels of E-cadherin in the newly formed junction (Fig. 5B,E,F), consistent with our findings that E-cadherin turnover is reduced in Ecad-LARIAT embryos, and with a model suggesting that E-cadherin planar polarity requires endocytotic trafficking [Levayer et al., 2011]. We conclude that the increase in cell adhesion seen in Ecad-LARIAT embryos has a dramatic impact on cell rearrangements despite a lack of significant changes in myosin levels or distribution.

**Figure 5.**
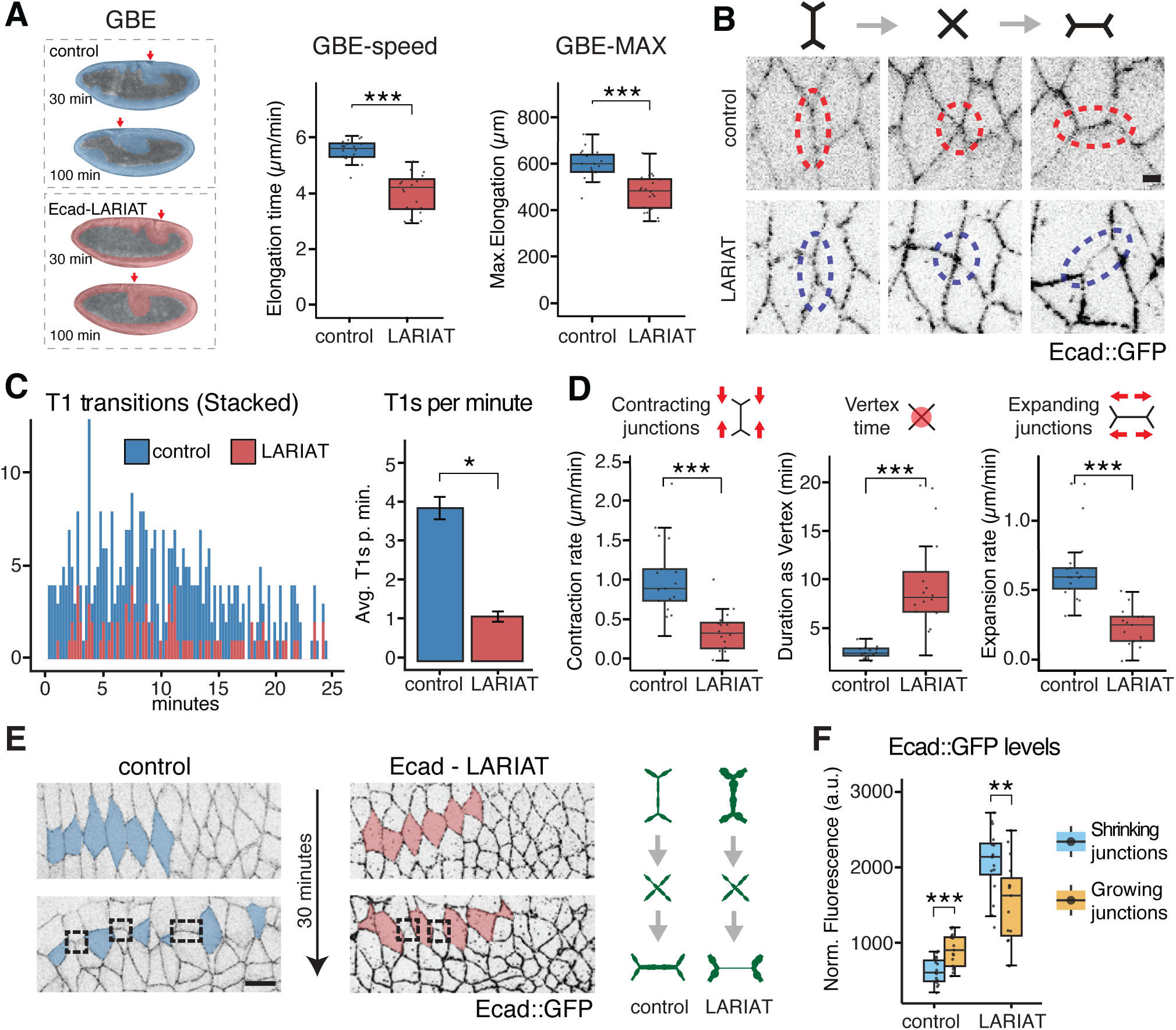
LARIAT-mediated E-cadherin clustering impairs cell rearrangements. **(A)** Brightfield images of control and Ecad-LARIAT embryos showing defects in germband extension (GBE; red arrows indicate tip of germband). The germband is false colored in red (LARIAT) and blue (control). Speed of GBE and maximal elongation are quantified and show delays in Ecad-LARIAT embryos compared to controls. Statistics: Wilcoxon rank-sum test, GBE speed p = 1.4 × 10^-9, GBE max p = 1.6 × 10^-6. Sample sizes: control = 20 embryos, LARIAT = 20 embryos. **(B)** T1 transitions in control and Ecad-LARIAT embryos. Stills from movies, showing a shrinking junction, a vertex, and a growing junction (left to right). Scale bar 2µm. **(C)** Number of T1 transitions is significantly reduced in Ecad-LARIAT embryos compared to controls. Stacked bar plot showing the number of T1 transitions per minute over the first 25 minutes of GBE (left) and average T1s per minute (right). Comparisons were assessed with a linear mixed-effects model, p = 0.0104, control = 3 embryos, LARIAT = 3 embryos. **(D)** T1-kinetics are significantly altered in Ecad-LARIAT embryos. Junction contraction rate (µm/min) is reduced, vertex resolution time (min) is increased, and junction expansion rate (µm/min) is reduced in Ecad-LARIAT embryos. Dots represent individual junctions. Shrink-rate comparisons were assessed using a linear mixed-effects model: contraction p = 2.8 × 10^-5, vertex p = 3.5 × 10^-6, expansion p = 1.8 × 10^-6. Sample sizes: control = 5 embryos, 17 junctions; LARIAT = 7 embryos, 16 junctions. **(E)** Cell arrangement in control and Ecad-LARIAT embryos before (top) and after 30 minutes of GBE showing that tissue elongation is impaired in Ecad-LARIAT embryos. Growing or newly formed junctions during cell intercalation are highlighted in dotted boxes. Illustration of E-cadherin levels in shrinking and growing junctions (right). Scale bar 10µm. **(F)** Quantification of E-cadherin levels at shrinking and growing junctions shows a relative increase in control and a significant decrease in E-cadherin levels in growing junctions in Ecad-LARIAT embryos. A linear mixed-effects model was used for statistical comparison: control comparison p = 4.3 × 10^-8, Ecad-LARIAT comparison p = 0.0036. Sample sizes: control = 5 embryos, 17 junctions; LARIAT = 7 embryos, 16 junctions.

### A vertex model with dissipative adhesion simulates reduced cell rearrangements during convergent extension

Quantitative understanding of how cell adhesion contributes to morphogenesis requires knowledge of tissue-specific rheological properties [Mongera et al., 2018; Petridou et al., 2021; Kim et al., 2021]. Because cells act collectively and are mechanically constrained by neighboring tissues, it is experimentally challenging to isolate and control all parameters influencing large-scale movements. Biophysical modeling may identify key parameters of complex processes by independently varying mechanical inputs and predicting emergent tissue-scale outcomes [Bi et al. 2015; Lawson-Keister and Manning, 2021]. In Drosophila, two-dimensional vertex models have been applied to describe epithelial packing and emergent collective behaviors [Rauzi et al., 2008; Wang et al., 2020].

In vertex models, tissue mechanics emerge from the collective behavior of individual epithelial cells, each represented as a polygon that moves its vertices to minimize a global energy function (Fig. 6A, Fig. S6A,B). The energy of each cell comprises an (i) area elasticity term, which penalizes deviations from a preferred area, (ii) a perimeter elasticity term, which captures the balance between cortical tension and adhesion, and (iii) a line tension term, which represents polarized edge contractility. Adhesion is incorporated through the preferred perimeter, with higher adhesion corresponding to larger preferred perimeters and reduced effective contractility [Honda and Eguchi 1980; Bi et al., 2015]. Consequently, tissues with high adhesion are predicted to contain more irregularly shaped cells (expressed as a higher cell shape index), exhibit lower effective line tension, and possess a reduced threshold for topological rearrangements, resulting in more fluid-like behavior (Fig. 6C). We tested predictions of this standard model by simulating an increase in adhesion (implemented as increased preferred cell perimeter) in the ventral ectoderm to represent the elevated adhesion in Ecad-LARIAT embryos. The model predicted a reduction in mean edge tension, enhanced cell rearrangements and convergent extension (Fig. 6C, Fig. S6C, Video S7), suggesting an increase in tissue fluidity.

**Figure 6.**
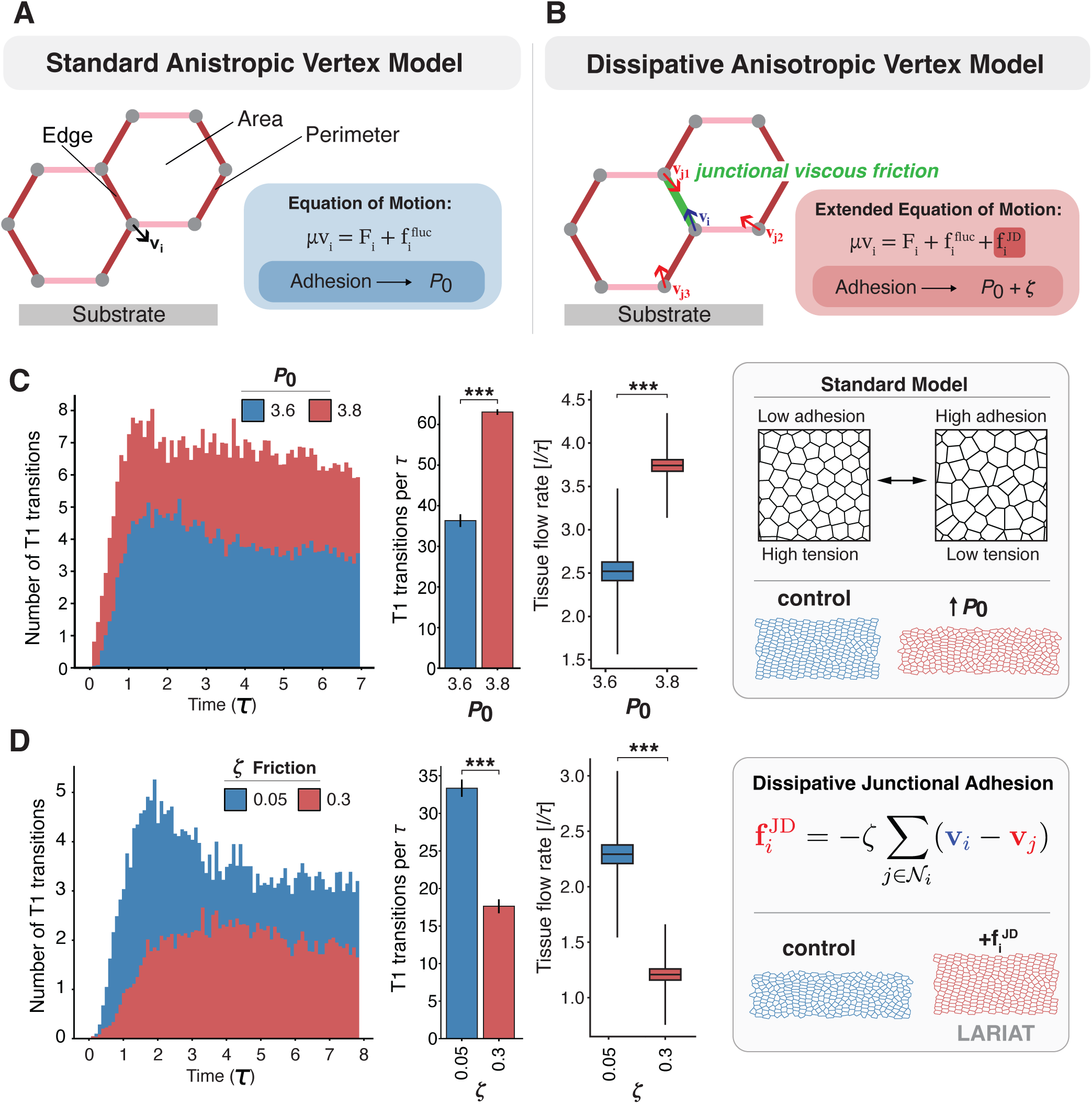
Increased adhesion friction, not cell shape changes, explains clustering-induced tissue rigidification. **(A)** Standard anisotropic vertex model. Two adjacent cells with a shared junction. Vertices of the cells (and cells as a result) move according to the equation of motion. *μ* is the substrate friction, **v**_*i*_ is the velocity of a vertex, **F**_*i*_ is the mechanical force acting on that vertex, which is defined by the mechanical energy *E* (see Methods, Fig. S6A,B), and 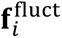 represents stochastic fluctuations. Mechanical energy includes area and perimeter elasticity terms as well as polarized tension. Darker edges indicate junctions with higher polarized tension. In standard models, adhesion is incorporated only through the preferred perimeter *P*_0_. **(B)** Dissipative anisotropic vertex model. Two adjacent cells showing viscous junctions (indicated by green line). Vertices move according to the extended equation of motion, where the additional friction term (**f**^()^) represents junction-specific friction from E-cadherin adhesion (see Methods, Fig. S6A,B). This friction term is proportional to the relative velocity (**v_i_ − v_j_**) between adjacent vertices. Other terms are the same as in the standard model. Darker edges indicate polarized tension. Adhesion is incorporated through the preferred perimeter *P*_0_ and the friction coefficient *ζ* that resists relative motion between adjacent vertices, capturing the frictional resistance from E-cadherin clustering. **(C)** Standard anisotropic vertex model (Video S7) predictions contradict experimental observations. Number of T1 transitions over time (left) and T1 transition rate per simulation time (middle) for two preferred shape index values: *P*_0_ = 3.6 (blue, baseline) and *P*_0_ = 3.8 (red, to account for increased adhesion in standard models). Tissue flow rate for standard vertex model (right). Higher shape index predicts faster convergent extension, opposite to the impaired extension observed with Ecad-LARIAT embryos (Fig. 5). Illustration on the right shows tissue geometry and tissue extension with different cell shapes (*P_0_*). **(D)** Dissipative anisotropic vertex model (Video S8) captures experimental phenotypes. Number of T1 transitions over time (left) and T1 transition rate (middle) for two values of junction friction coefficient: *ζ*=0.05 (blue, control/wild-type) and *ζ* = 0.3 (red, clustering/Ecad-LARIAT). Preferred shape index (*P*_0_ = 3.6) is fixed for both conditions as experimental measurements of cell shape index showed no significant difference between control and Ecad-LARIAT at t=0, (control: 3.99 ± 0.02, (n=4 embryos); LARIAT: 4.00 ± 0.03, (n=3 embryos); p=0.63, Mann-Whitney U test). Increasing junction viscosity substantially reduces T1 transition rates and tissue flow rate in the dissipative model. Increased junction friction dramatically slows convergent extension, consistent with experimental measurements showing impaired germband elongation in Ecad-LARIAT embryos (Fig. 5), which the standard models cannot explain. Illustration on the right shows how dissipative junctional adhesion is defined and tissue extension with different friction regimes. **All simulations:** n = 80 simulations per condition. Bar plots (C-D, middle) show mean and SEM Box plots (C-D, right) show mean (central line), box edges extend to ±1 SEM, and whiskers extend to ±1 SD. Time units in *τ* and length units in *l*. Other parameters: *A*_0_ = 1, *K*_*A*_, = 1, *K*_*P*_ = 1, *μ* = 1 and T=0.02 in natural simulation units. Polarized tension was reduced from *γ*_0_=0.3 (control) to *γ*_0_ = 0.28 (Ecad-LARIAT) to reflect the minimal decrease in mean polarized Myosin intensity (Fig 4D). *** Mann-Whitney U test, p < 0.001.

However, this contradicts our experimental observations (Fig. 5), which suggest that E-cadherin clustering/stabilization and enhanced cell adhesion interferes with cell rearrangements causing a dramatic decrease of T1 transitions. To reconcile this discrepancy, we hypothesized that clustered E-cadherin complexes create frictional resistance to junctional remodeling, due to lack of E-cadherin mobility. To test this, we introduce an additional term into the vertex model representing *dissipative cell adhesion* (*ζ*; or cell-cell friction) (Figs. 6B and S6B, see Methods) in the equation of motion. Previous work has incorporated internal dissipation arising from cell-cell interactions [Tong et al. 2022, Rozman et al. 2025], but the effect of enhanced dissipation from increased adhesion has not been studied in the context of convergent extension. Here we apply this framework to an anisotropic vertex model where tissue flows arise from oriented cell rearrangements. Incorporating this term allowed edge tension to reduce minimally (Fig. S6D) while substantially increasing resistance to cell rearrangements (Fig. 6D). Simulations under these conditions predict pronounced slowing of morphogenetic flows during germband extension (Fig. 6D; Video S8), consistent with our experimental data of Ecad-LARIAT embryos.

### E-cadherin clustering interferes with cell rearrangement but not apical constriction

A modified vertex model that includes a dissipative cell adhesion term and our analysis of cell intercalation during germband extension in Ecad-LARIAT embryos predicts that morphogenetic movements that require cell-cell contact changes as cells slide along each other are impacted by enhanced E-cadherin clustering due to an increased energy barrier. To further test this prediction, we compared two movements that rely on apical constriction. The apical constriction of ingressing neuroblasts is accompanied by cell-contact changes between the neuroblasts and non-ingressing cells and between adjacent non-ingressing cells as neuroblasts delaminate from the epithelium [Simoes et al., 2017]. In contrast, the invagination of the mesoderm is executed by the meta-synchronous apical constriction of a coherent field of cells which, however, do undergo little to no junctional cell rearrangement during invagination [Perez-Vale and Peifer, 2020; Ko and Martin 2020].

Mesodermal apical constriction is driven by pulsatile accumulations of medial myosin, producing rapid apical surface reduction across a continuous epithelial monolayer [Martin et al., 2009]. The loss of the CCC disrupts cell adhesion between mesodermal cells and compromises invagination [Sawyer et al., 2009; Martin et al., 2010; Sheppard et al., 2023]. Apical constriction of mesodermal cells appeared normal in Ecad-LARIAT embryos (Fig. 7A; Video S9), suggesting that constriction itself, when it proceeds without neighbor exchange, is insensitive to E-cadherin stabilization. These observations suggest that cell adhesion strength does not have a marked impact on cell constriction-based invagination in the absence of cell rearrangement.

**Figure 7.**
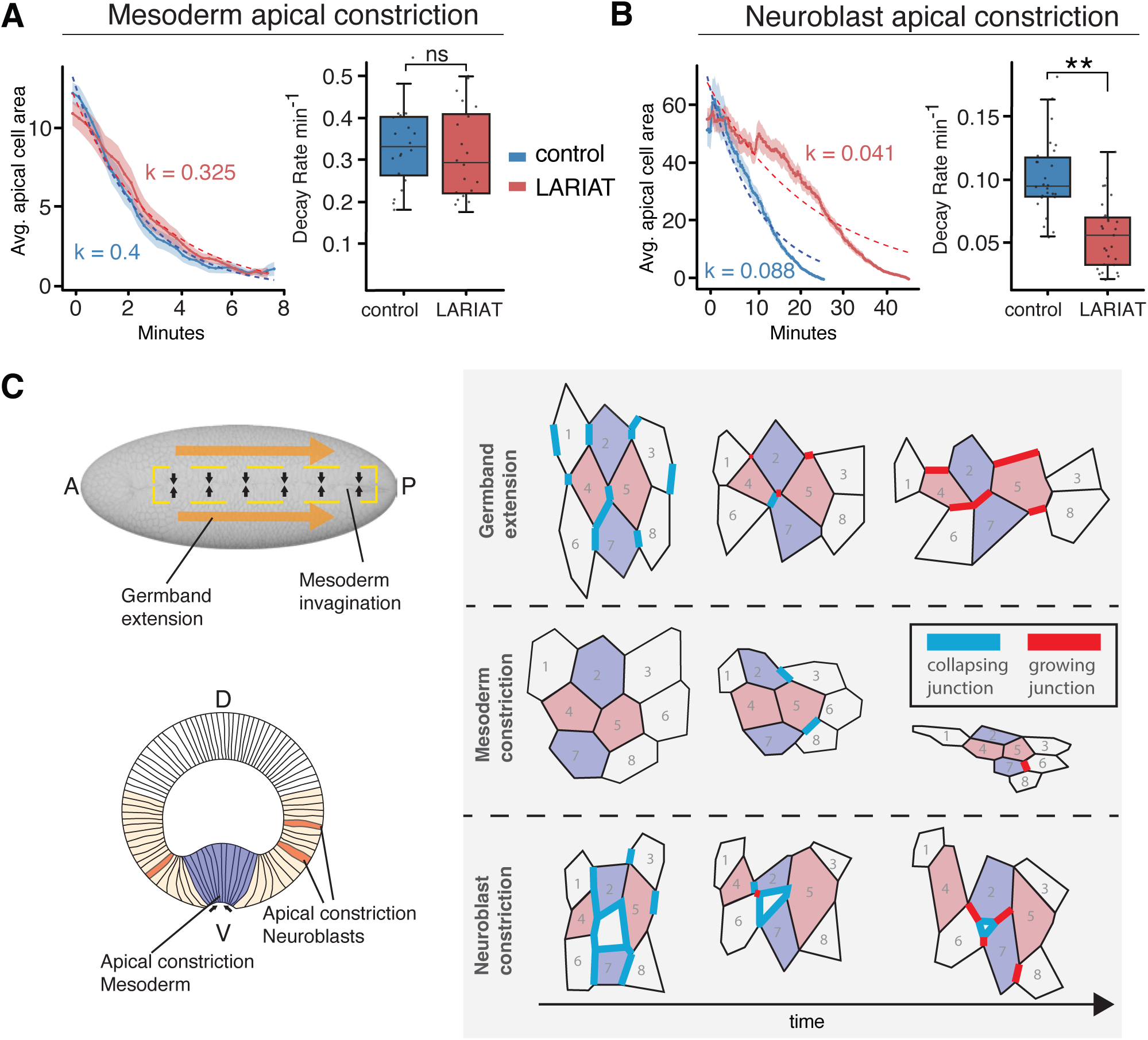
Clustering of E-cadherin differentially affects morphogenetic movements. **(A)** Mesoderm apical constriction, quantified as a decay rate of apical area over time, is not affected in Ecad-LARIAT embryos. Dots represent individual cells. Statistics: Comparisons were assessed with a linear mixed-effects model, mesoderm p = ns, control: 4 embryos, 20 cells, LARIAT: 4 embryos, 20 cells. **(B)** Neuroblast apical constriction is delayed in Ecad-LARIAT embryos. Statistics: Comparisons were assessed with a linear mixed-effects model, p = 0.00235, control: 5 embryos, 26 cells, LARIAT: 7 embryos, 31 cells. **(C)** Illustration of three morphogenetic movements during Drosophila gastrulation (left). Traces from live embryos showing cell contact changes during germband extension, mesoderm invagination, and neuroblast ingression. Germband extension and neuroblast ingression involves the shrinkage and collapse of cell-cell contacts and the formation of new junctions illustrated in blue and red, respectively. In contrast, of the invagination of the mesoderm relies on apical constriction without significant cell contact changes.

In contrast, neuroblast ingression, a developmental epithelial-mesenchymal transition that depends on junctional remodeling and endocytic removal of apical membrane [Simões et al. 2017; 2022; An et al. 2017], was significantly delayed in Ecad-LARIAT embryos (Fig. 7B; Video S10). The kinetics of apical area loss was reduced with a lower decay rate of apical area and an average delay of ∼15 minutes (∼107%) in the time period a neuroblast requires for ingression (Fig. 7B). Neuroblast constriction relies on anisotropic edge contraction and junctional turnover at interfaces between neuroblasts and non-ingressing cells and between adjacent non-ingressing cells [Simoes et al., 2017] similar to the edge contractions during T1 transitions. We conclude that the reduced mobility of E-cadherin likely impedes remodeling dynamics, increasing the yield stress required for cell-contact contraction and lowering effective actomyosin-driven apical constriction. These findings support the idea that dynamic adhesion is required for efficient active constriction when it involves cell-cell contact changes in the plane of the epithelium (Fig. 7C).

## Discussion

Analyzing the contribution of E-cadherin and cell adhesion to morphogenetic movements within epithelial layers has been challenging. Loss-of-function of E-cadherin or associated core components of the CCC causes a loss of epithelial tissue architecture interfering with investigating the role of E-cadherin in specific epithelial cell movementst, such as the cell intercalations that drive convergent extension of the Drosophila germband. In contrast to the pleiotropic effects of E-cadherin loss-of-function, E-cadherin overexpression has little to no effect on morphogenesis in Drosophila [Pacquelet and Rorth, 2005; Wang et al., 2024; this work] or vertebrate models [e.g. Ninomiya et al., 2012]. When overexpressed, excess E-cadherin is efficiently removed from the surface, suggesting a robust intrinsic control of cell adhesion during development. Here we took advantage of E-cadherin clustering as an alternative approach to generating an E-cadherin adhesion gain-of-function. In vitro studies have identified clustering of the CCC as a key determinant of adhesion strength [Hong et al., 2010; Strale et al., 2015; Bertocchi et al., 2017; Slováková et al., 2022; Yap et al., 2015; Ishiyama & Mège, 2017; Troyanovsky, 2023; Monster et al., 2025; Troyanovski et al., 2025]. It is generally thought that E-cadherin clusters are also the principle components of supramolecular adhesion complexes in vivo [e.g. Quang et al., 2013]. However, how clustering contributes to morphogenesis remained largely unexplored. We found that enhanced clustering using LARIAT or the CRY2 component of the LARIAT system has dramatic effects on E-cadherin dynamics including increased surface levels and cluster size, reduced lateral mobility, and reduced protein turnover. These parameters are consistent with a substantial increase in cell adhesion without disrupting epithelial tissue integrity and provide a new strategy for manipulating cell adhesion in vivo.

Ecad-LARIAT embryos showed enhanced recruited of β-Catenin and α-Catenin to E-cadherin clusters but did not affect the junctional distribution or levels of actin or myosin. Interestingly, we found that Vinculin is reduced at AJs in Ecad-LARIAT embryos. Vinculin is recruited to AJs by binding to the mechanosensitve M-region of α-Catenin [Yonemura et al., 2010; Kale et al., 2018; Sarpal et al., 2019; Sheppard et al., 2023]. The reduced recruitment of Vinculin is consistent with the distribution of tension forces – which appear to be similar in control and Ecad-LARIAT embryos – over a larger number of CCC complexes thereby reducing the force applied to individual adhesion complexes. Collectively, these findings suggest that the E-cadherin clusters in Ecad-LARIAT embryos assemble a normal CCC that is responsive to mechanical forces.

E-cadherin in Ecad-LARIAT embryos showed a uniform distribution at horizontal and vertical junctions in the germband in contrast to wild-type where E-cadherin is somewhat enriched at horizontal junctions [Blankenship et al., 2006; Rauzi et al., 2008; Levayer et al., 2011]. Myosin, in contrast, remained enriched at vertical junctions. It was proposed that the E-cadherin reduction at vertical junctions is the result of myosin contraction and activation of the formin Diaphanous, fostering clustering and hence endocytotic removal [Levayer et al., 2011; Levayer and Lecuit, 2013; Quang et al., 2013]. Our data suggest that increasing cluster size is not an obligatory trigger for the endocytotic removal of E-cadherin, pointing to more complex mechanisms that regulate CCC cluster size and endocytosis. E-cadherin in Ecad-LARIAT embryos showed a much-reduced turnover which may account for the lack of asymmetry in E-cadherin distribution.

Most T1 transitions are blocked in Ecad-LARIAT embryos. The remaining T1 transitions are slowed at each step of the process: contraction of vertical contacts, dwell time at four cell contacts, and formation of new horizontal junctions. Interestingly, new horizontal junctions accumulated only a small amount of E-cadherin compared to wild-type controls. This is consistent with a transcytosis model where E-cadherin is removed from shrinking vertical junctions and used to establish the new horizontal junctions. The finding that E-cadherin planar polarity seemed lost, while myosin planar polarity was retained in Ecad-LARIAT embryos, suggests that contractile mechanisms driving germband extension are in place, yet insufficient to overcome increased adhesion.

E-cadherin clustering and turnover at AJs sets the energy barrier for cell rearrangements. By enhancing E-cadherin clusters, we increased this barrier and consequently the yield stress that would be required to overcome this resistance. The stabilization of junctions is apparently not overcome by enhanced actomyosin force generation as both junctional actin and myosin levels and recoil after laser cutting of AJs remained close to controls. Consequently, cell rearrangements are blocked or slowed, reducing fluidity and tissue flow. This is inconsistent with previous models of tissue mechanics that were based on cell shape and suggested that enhanced cell adhesion would promote fluidity and cell rearrangements, [Bi et al., 2015; Wang et al., 2020; Lenne and Trivedi, 2022]. Unlike standard vertex models where all dissipation comes from medium drag and adhesion is coupled with contractility in the perimeter term [Bi et al., 2015; Wang et al., 2020], our dissipative junctional anisotropic vertex model includes explicit cell-cell friction through junction-specific viscous forces. These viscous forces can independently regulate tissue dynamics by increasing resistance to cell motion rather than by changing cell shape, consistent with our in vivo observations. The junctional viscosity coefficient represents effective friction from E-cadherin-mediated adhesion. While recent work uses internal dissipation to study nematic-activity-driven flows [Rozman et al. 2025], we focus on how this friction affects contractility-driven convergent extension, where tissue flow arises from polarized myosin activity rather than nematic stresses. Higher junction viscosity in this regime reduces both T1 transition rates and overall tissue flow. Recent detailed modeling of apposed cortices [Nestor-Bergmann et al. 2022] predicts that high adhesion friction promotes vertex jamming and unresolved rosette formation, which is in line with T1 rates we observe, especially increased vertex times. While our coarse-grained approach enables modeling tissue-scale flow, it cannot capture the detailed microscale mechanics of apposed cortices or individual adhesion bond dynamics. However, we introduce a term that captures the net consequence of strong adhesion for tissue-scale rearrangement dynamics, successfully modeling the consequences of increased adhesion on tissue morphogenesis.

Adhesion is deployed differently across morphogenetic processes, allowing embryos to tune tissue mechanics to the specific demands of each movement. In mesoderm invagination, where collective apical constriction without cell-cell contact change is the dominant driver [Martin et al., 2009], apical constriction and invagination appeared normal in Ecad-LARIAT embryos. By contrast, during neuroblast ingression, apical constriction is coupled to progressive junctional remodeling and endocytic removal of apical membrane [Simoes et al., 2017, 2022]. The cell-cell contact changes between ingressing neuroblasts and non-ingressing cells and between adjacent non-ingressing cells require cadherin turnover that is apparently inhibited through LARIAT-mediated clustering of E-cadherin, significantly delaying neuroblast ingression. The communalities between neuroblast ingression and cell intercalation during convergent extension of the germband is the requirement for cell-cell contact change that drives tissue remodeling in contrast to mesoderm invagination where this is not the case (Fig. 7C).

Our findings are consistent with the view that adhesion does not simply promote or oppose morphogenesis or maintain tissue integrity while other forces conduct tissue remodeling but instead helps to define the viscoelastic properties of cells and tissues, including how readily they deform, dissipate stress, or remodel contacts under force. The modulation of expression levels of E-cadherin has apparently little impact on the regulation of adhesion force and thus morphogenesis as cells effectively control E-cadherin surface levels. On the contrary, we propose that the regulation of E-cadherin clustering plays a major role in controlling friction between cells that tune the speed of cell-cell contact changes and tissue remodeling in the presence of constant actomyosin force generation. Clustering of E-cadherin and the CCC can be regulated by multiple mechanisms including the cis- and trans-interactions of the cadherin extracellular domains and the association of the CCC with the actin cytoskeleton that is principally mediated by α-Catenin and its multiple actin-associated binding partners [Yap et al., 2015; Strale et al., 2015; Chen et al., 2015; Trojanovsky, 2023]. Cadherin extracellular domain induced clustering and actin-binding mediated clustering act independently and can synergize to enhance clustering [Trojanovsky et al., 2025]. How clustering of the CCC is regulated in specific cell and tissue movements remains to be explored. The interplay between adhesion generated friction and force generation by actomyosin or other processes determine the dynamics of tissue movements that require cell-cell contact changes. This interplay could therefore make key contributions to developmental timing and morphological change during evolution.

## Acknowledgements

We thank Xiaobo Wang, Eurico Morais-de-Sa, Adam Martin, Yohanns Bellaiche, Jennifer Zallen, Hiroki Oda, Thomas Lecuit, Daniel St. Johnston, Tony Harris, Rodrigo Fernandez-Gonzalez, Gertrude Schüpbach, the Bloomington Drosophila Stock Center, the Drosophila RNAi Screening Center at Harvard Medical School, and the Developmental Studies Hybridoma Bank for fly stocks and reagents. We thank Rodrigo Fernandez-Gonzalez for advice on data analysis and we appreciate critical reading of the manuscript by Dorothea Godt. We would like to thank Imaging Facility of the Department of Cell and Systems Biology and the Imaging Facility of Hospital of Sick Children, University of Toronto, for support.

## Author contributions

G.L., G.E.-T., and U.T. developed the project. G.L. carried out most of the experimental work. Additional experimental work was conducted by S.S. and F.Z. The biophysical model was developed and implemented by E.E. and G.E.-T., G.L., G.E.-T., and U.T. wrote the paper. G.E.-T. and UT raised funds to support the project.

## Data and material availability

All data are available in the main text or the supplementary materials.

## Supplementary materials

Methods

Supplementary Figures S1 to S6

Supplementary Tables S1

Supplementary Videos S1 to S10

References associated with supplementary materials.

