## Supplemental Materials for "E-cadherin clustering as a regulator of morphogenesis"

April 17, 2026

### Supplemental materials


**Authors**

Gerald Lerchbaumer[1], Sergio Simoes[1], Ermia Etemadi[2], Fadi Zidan[1], Gonca Erdemci-Tandogan[2,3] and Ulrich Tepass[1]*

**Affiliation**

(1) Department of Cell and Systems Biology, University of Toronto; Toronto, ON M5S 3G5, Canada

(2) Department of Physics and Astronomy, Western University, London, ON, Canada

(3) Department of Medical Biophysics, Western University, London, ON, Canada

**Correspondence should be addressed to:**

Ulrich Tepass
ORCID: 0000-0002-7494-5942
Department of Cell and Systems Biology
University of Toronto
25 Harbord Street
Toronto, Ontario M5S 2M6
Canada


**Supplemental materials included:**

Methods

Supplemental Figures S1-S6 plus legends

Supplemental Tables S1

Supplemental Videos S1 – S10

### Methods

Reagents and fly lines used in this study are listed in Table S1.

#### Drosophila genetics

**E-cadherin overexpression:** To increase levels of E-cadherin expression we crossed *ubi-Ecad::GFP* (Oda and Tsukita, 2001), *matαtub67-Gal4*; *sqh-Sqh::mCherry* flies to *UASp-shg::GFP* (BDSC 48445) flies and analyzed the F2 progeny.

**Optogenetics:** For E-cadherin optogenetic clustering *matαtub67-Gal4, endo-Ecad::GFP*; *sqh-Sqh::mCherry* flies were crossed to *endo-Ecad::GFP; UASp-LARIAT/TM6B* (gift from X. Wang). The F1 progeny was sorted against *TM6B* and crossed in the dark at 25°C. F2 embryos were subsequently imaged as described below. As a control F1 progeny flies containing *TM6B* instead of *UASp-LARIAT* were used. For optogenetic clustering of PH::GFP we crossed *matαtub67-Gal4*, *UAS-PH::GFP* (Gervais et al., 2008) to *UAS-PH::mCherry*; *UASp-LARIAT* flies. For optogenetic Ecad-CRY2 clustering we crossed *endo-Ecad::GFP, matαtub67-Gal4; sqh-Sqh::mCherry* flies to *endo-Ecad::GFP; UASp-CRY2vhh/TM6B* flies. For live imaging of α-Catenin during optogenetic clustering of E-cadherin, we generated a line carrying *endo-Ecad::GFP, matαtub67-Gal4; αCat::KRFP* and crossed it to *endo-Ecad::GFP; UASp-LARIAT* flies. For live imaging of *Vinc::mCherry* during optogenetic clustering of E-cadherin, we generated a line carrying *endo-Ecad::GFP, Vinc::mCherry* (Kale et al., 2018); *UASp-LARIAT* and crossed it to *matαtub67-Gal4*, *endo-Ecad::GFP* flies. F2 progeny of all crosses above were raised in the dark and imaged as described below.

#### Live imaging

For imaging, embryos were dechorionated in 50% bleach and mounted in halocarbon oil 27 (Sigma-Aldrich) between an oxygen permeable membrane (YSI; Xylem Inc.) and a coverslip (no. 1.5).

For **confocal imaging**, fluorophores were excited using an optically pumped semiconductor laser (OPSL) at 488nm for GFP and 552nm for mCherry. Laser intensities varied from 2-5% depending on which fluorophore was used. A Leica SP8 was used for confocal imaging. The following objective lenses were used: Leica HCX Plan-Apochromat 63x/1.4 NA CS2, Leica HCX Plan-Apochromat 40x/1.3 NA CS2. HyD detectors (Gain 100%) were used for all live imaging. For time lapse movies with two fluorophores, sequentially imaging between lines and bidirectional scanning was used. For live imaging of germband extension, neuroblast ingression, and mesoderm invagination the ventral ectoderm was imaged at a depth of 7 µm (15 steps at 0.5 µm per optical section). A stack was taken every 15 seconds if not stated otherwise in the figure legend. Pixel dimensions were 180nm/pixel. Confocal imaging: Fig. 1C; Fig. 2A,E; Fig. 3 A-D; Fig. 4A,C,D; Fig. 5 B-F; Fig. 7A-B; Fig. S1 A,B; Fig. S2B,C; Fig. S3D; Fig. S4A-C; Fig. S5A,C.

For **spinning disk microscopy**, embryos were dechorionated and mounted as mentioned above and imaged with a Nikon Ti2 inverted microscope equipped with a CSU-X Spinning Disk (Yokogawa), a sCMOS camera (Teledyne Photometrics Prime 95B), a 100X Plan Apochromat VC NA 1.4 oil-immersion objective (Nikon) and a 60X Plan Apochromat lambda NA 1.4 oil-immersion objective (Nikon). Spinning disc imaging: Fig. 2D; Fig. 4F; Fig. S3C.

For **bright-field microscopy** a Carl Zeiss Axiophot2 microscope using a phase-contrast 20X lens (NA 0.5) or a 10X lens paired with a Canon Rebel XSi digital camera was used for embryonic cuticles and time lapse movies of whole embryos for germband extension analysis. Light microscopy: Fig. 5A.

Movies from confocal and spinning disk microscopy were acquired as 4-dimensional stacks and flattened by maximum intensity projection in ImageJ.

For **SIM-imaging** a commercial structured-illumination microscope (MI-SIM, CSR Biotech) was used to acquire and reconstruct Ecad::GFP images in the germband (LARIAT and control embryos) with a 100x objective (NA = 1.5). Imaging followed the protocol of [Huang et al., 2018]. To further enhance resolution and contrast, sparse deconvolution was applied [Huang et al., 2018, Zhao et al., 2022]. SIM-imaging: Fig. 1D-G, Fig. S1B.

**Optogenetic clustering of E-cadherin**

Homozygous flies for endogenously tagged E-cadherin, carrying a maternal driver and one copy of *UASp-LARIAT* or *UASp-CRY2vhh*, as well as one copy of *Sqh::mCherry* (described in fly genetics) were raised in the dark and dechorionated under light with a high-pass filter and kept in the dark until imaging. Embryos were oriented under the confocal microscope using a RFP filter. Imaging was performed as described in the live imaging sections. Photo-activation was achieved by excitation with an OPSL at 488nm wavelength at 3% power. This approach led to activation of the LARIAT system in the whole embryo which was subsequently imaged continuously.

**Fluorescence recovery after photobleaching (FRAP)**

FRAP experiments were performed in two modes: whole-junction FRAP and spot-junction FRAP, each using distinct acquisition and analysis pipelines.

**Whole-junction FRAP:** Stage 7–8 embryos were imaged on a Leica SP8 confocal microscope using LASX software (version 5.3.1). Entire bicellular AJs were photobleached with the 488 nm laser at 60% intensity, and fluorescence recovery was recorded in $z$-stacks spanning 7µm at 15s intervals in two channels (488 = Ecad::GFP, 552 = Sqh::mCherry). For each experiment, raw fluorescence intensities of the bleached junction were measured in Pyjamas [Fernandez-Gonzalez et. al., 2022], and recovery was measured by normalizing mode-subtracted raw fluorescence intensity to an unbleached reference junction in proximity, which served as an internal control for acquisition variability and overall photobleaching. The reference-corrected traces were then rescaled by subtracting the first post-bleach value and dividing by the difference between the pre-bleach and post-bleach values, such that the first post-bleach frame was set to 0 and the pre-bleach level to 1.

**Spot-junction FRAP.** Stage 7–8 embryos (Control, LARIAT, and E-cadherin overexpression) were imaged on a Nikon Eclipse Ti2-E Yokogawa CSU-X1 spinning disk with a 60X oil immersion objective (N.A. 1.4). Images were acquired at 12-bit depth (pixel size 0.18 µm) with 600ms exposure, looping three times pre-bleach and 40 times post-bleach at 15 s intervals. Circular ROIs of radius 0.72 µm were photobleached using 50% 488nm laser power for 1s, reducing junctional Ecad::GFP intensity by 78–99%. Background and junction ROI intensities were measured manually in Fiji, with drift correction in Pyjamas and local reference regions (200×200px) used to normalize acquisition variability. Corrected and normalized FRAP intensities were computed as above.

**Model fitting**.

Recovery curves were then fit to a one-phase exponential model

$$F(t) = F_0 + (1 - F_0)(1 - e^{-kt}),$$

yielding recovery rate constants ($k$), and half-times ($t_{1/2} = \ln(2)/k$).

Mobile fractions were not derived from the fit, but instead calculated directly from the experimental recovery traces as:

$$\text{Mobile Fraction} = \frac{F_{\text{end}} - F_0}{F_{\text{prebleach}} - F_0},$$

where $F_{\text{end}}$ is the mean normalized intensity at the final three post-bleach time points, $F_0$ is the intensity immediately after bleaching, and $F_{\text{prebleach}}$ is the mean pre-bleach intensity. Fits were performed in R using nlsLM (minpack.lm). Statistical comparisons between two groups were performed using Wilcoxon rank-sum test (Mann–Whitney) and a Kruskal–Wallis with Dunn's test (Bonferroni corrected) for three groups due to non-normal distribution of data. Data are displayed as mean $\pm$ SEM with significance annotated (ns, *, **, ***).

**Laser ablation**

Laser ablation experiments were performed to measure junctional recoil behavior and were done with a Optimicroscan UV galvo scanner (Nikon) using a 355 nm laser at 1,000 Hz, with a 200 s pixel dwell time, and a 4 µm line of ablation. Ablations were done using a 60X Plan Apochromat lambda NA 1.4 oil-immersion objective (Nikon). Subsequent imaging was done with a spinning disk Nikon microscope as described above. Vertical (DV oriented) junctions were cut in stage 7-8 embryos. Junction distances were tracked over time post-ablation with a recoil velocity plugin in Pyjamas [Fernandez-Gonzalez et. al., 2022], and data were recorded at 3 second intervals in a 3 µm stack with a 0.3 µm interval.

Average recoil distance curves were generated by calculating mean and standard error of distances at each time point. Maximum recoil distances were extracted for each junction, and statistical comparisons were performed using Wilcoxon rank-sum tests to evaluate differences between control and LARIAT treatments. Initial recoil velocities were also calculated for each junction, statistically compared via Wilcoxon rank-sum tests, and visualized using boxplots with jittered points.

**Immunohistochemistry**

Embryos were fixed in 4% formaldehyde in PBS (pH 7.4) and heptane for 20 minutes. Devitillinization was performed in a 1:1 methanol/heptane mix by agitation. The following antibodies were used: anti-GFP (polyclonal Alexa fluor 488, Invitrogen, 1:400), anti-Arm (mouse monoclonal N2 7A1, 1:100; DSHB), Phalloidin Alexa 647 (Thermo Fisher), anti-aCat (p121; 1:1000, this lab), anti-Crb [Pellikka et al., 2002]. Secondary antibodies conjugated to Alexa Fluor 488, Alexa Fluor 568, or Alexa Fluor 647 (Invitrogen) were used at 1:400. Embryos were mounted in Vectashield (ThermoFisher) and imaged using a confocal microscope as described above.

**Quantification of E-cadherin cluster size**

MI-SIM $z$-stacks encompassing the AJs (16-bit images, voxel size $32.5 \times 32.5 \times 180$ nm in $x, y, z$) were thresholded in FIJI using the *Intermodes* method. The resulting binary stacks were analysed with FIJI's *3D Objects Counter* plugin to obtain cluster volumes; objects smaller than $1 \times 10^{-3}\ \mu\mathrm{m}^3$ were excluded from further analysis.

**Colocalization analysis**

Colocalization analysis between Ecad::GFP and αCat, Arm, Vinc or F-actin was done by choosing a single Z junctional plane in embryos co-stained for Ecad::GFP and αCat, Arm or F-actin (fixed samples) or co-expressing Ecad::GFP and Vinc::mCherry (live samples). All images used for colocalization were acquired with identical laser power, detector gain and offset settings within each experiment. A mild Gaussian blur ($\sigma = 0.5$ pixels; 1 pixel = 90 nm) was applied to reduce noise without affecting junctional structure width. A junctional binary mask was created in FIJI by manually drawing 7 pixel wide segmented lines over cell-cell junctions, using the Ecad::GFP channel, in both controls and LARIAT embryos. FIJI's plugin Coloc 2 was then used to calculate Spearman's rank coefficient between E-cadherin and the mentioned markers. No automatic or manual intensity thresholding was applied; all raw pixels within the junctional masks were included in the analysis. Negative controls were obtained by rotating the αCat channel in control or LARIAT embryos by 90 degrees clockwise and calculating the Spearman coefficient with the non-rotated Ecad::GFP channel on its correspondent junctional mask.

**Quantification of Ecad::GFP cluster density**

MI-SIM z-stacks encompassing adherens junctions were first converted into maximum-intensity projections in FIJI. Individual junctions were manually traced using the Segmented Line tool with a line width of 9 pixels (1 pixel = 0.0325 μm × 0.0325 μm). A custom FIJI macro was used to straighten each traced junction using the Straighten function. E-cadherin clusters were subsequently identified on the straightened junctions using Find Maxima with a peak-finding Prominence threshold of 15. Cluster density was calculated as the number of detected fluorescence maxima divided by the physical length of the corresponding straightened junction.

**E-cadherin cluster mobility analysis**

Spinning disc microscopy was used to analyze E-cadherin clusters at a time interval of 2.6 s. Cluster mobility was quantified by analyzing data obtained from ImageJ-generated kymographs (Reslice function), which were registered to a fixed tricellular junction reference point in Pyjamas [Fernandez-Gonzalez et al., 2022]. Individual E-cadherin clusters were chosen from stage 7 embryos based on visibility and tracked across time points. Clusters were tracked using the MTrackJ plugin in ImageJ. X and Y displacement over time was computed to quantify cluster mobility. Maximum displacement was shown for each cell by plotting X-coordinate maxima for each cluster. Displacement velocities were then calculated for each cluster as the change in X-position over time, statistically compared using Wilcoxon rank-sum tests, and visualized via boxplots. Subsequently, cumulative displacement over the entire measurement period was computed for each cluster by summing incremental displacements, visualized through boxplots and jittered data points. Total displacement was calculated as the difference between the maximum and minimum x-positions of each cluster. Velocity was determined as the frame-to-frame displacement over time. Cumulative displacement was computed as the sum of all positional changes along the x-axis.

**Planar polarity quantification**

For planar polarity analysis, images were obtained from timelapse movies at 10 min post germband extension onset, captured using a confocal microscope with a 63X objective as described above. An 80x50 micron region of interest (ROI) was selected on one side of the ventral midline in the central region of each embryo. Images were segmented using Tissue Analyzer (TA) and Epyseg software [Aigouy et al., 2016] to generate an analysis file containing junctional information used for subsequent analyses in R.

Junction orientation data from these segmented images were binned into categories: junctions between 75 deg. and 105 deg, relative to the ventral midline were classified as vertical, and junctions within 15 deg. of either 0 deg. or 180 deg. were considered horizontal. Intermediate orientations were excluded from horizontal-to-vertical ratio analyses. Fluorescence intensities for both E-cadherin and myosin II were background corrected by subtracting the ROI mode and averaged for each orientation bin, and the horizontal-to-vertical intensity ratios were calculated and statistically compared using Wilcoxon rank-sum tests. Individual dots in boxplot represent ratio of horizontal to vertical or vertical to horizontal ratio per cell.

**Particle Image Velocimetry (PIV) of medial myosin II flows**

Dual color movies of LARIAT and control early stage 7 embryos expressing Ecad::GFP and Sqh::mCherry were generated on a Leica SP8 Resonant Laser Scanner microscope, with resonant mode on. Nine z optical sections (0.895 um) separated by 0.5 mm were acquired with a 63X oil objective, NA = 1.4, every 5 seconds, for 10 minutes, at xy pixel size 144 nm. Hyperstacks were created by Z maximum projection in FIJI. To eliminate the effect of tissue and cell drift, individual cells were registered in Pyjamas [Fernandez-Gonzalez et al., 2021] by using the “Register image” function after assigning one fiducial to each vertex of the cell of interest in 40 frames (total duration: 200 sec). Single registered cells were then cropped in the Sqh::mCherry channel to an area of 120 x 120 pixels for PIV analysis. Images were further processed in FIJI with auto-level, Gaussian Blur with Sigma Radius 0.5, and brightness and contrast adjusted to set pixel min and max intensities equally in all movies, with the goal of enhancing medial myosin signal and removing background.

PIV was performed on single cells with PIVlab version 3.09 (Matlab) [Thielicke, 2014, 2021]. Frames were pre-processed via CLAHE (window size 64 pixels) and auto-contrast stretch (default). Each image (120 pixels X 120 pixels) was sub-divided into tiles during four iterative passes used for cross-correlation with the next frame and to obtain medial myosin II displacement in the x and y axes. Pass 1 consisted of 20 X 20 pixels tiles (= 2.88 X 2.88 mm) with 75% overlap; pass 2: 16 X 16 pixels tiles (= 2.3 X 2.3 mm) with 50% overlap; pass 3: 12 X 12 pixels tiles (= 1.73 X 1.73 mm) with 50% overlap and pass 4: 10 pixels tiles (= 1.44 X 1.44 mm) with 50% overlap, so that vectors in the vector field at each frame were separated by 5 pixels (= 0.72 mm). The PIV algorithm used for cross-correlation was Multi-pass FFT deformation, sub-pixel estimator 2 X 3 point and "extreme" correlation robustness. PIV vectors were then velocity-validated by applying a local median filter of 3 (threshold). No interpolation of missing vectors was used, to restrict our analysis only to observed medial myosin II clusters. To extract medial myosin displacement vectors, a polygonal ROI was manually drawn inside each cell, while avoiding junctional myosin signal in all time points. PIV vector magnitudes (mean speed, mean u and v vector components and |u|/|v| ratios) were extracted from that ROI at each time point (named "instantaneous"). The final vector fields shown were obtained by temporal integration (sum) of the respective displacement vectors during 25 sec (5 consecutive frames).

**Cell tracking and displacement analysis**

Cell displacement during germband extension (GBE) was quantified by tracking individual cells in time-lapse imaging datasets. Cell positions were extracted from registered movies using automated tracking, yielding x–y coordinates for each cell over time. For each track, positions were normalized by subtracting the initial coordinate, aligning all trajectories to a common origin to allow direct comparison of displacement across conditions. To visualize tissue flow, individual trajectories were plotted together with the mean trajectory per condition, calculated by averaging positions across cells at each relative timepoint. To quantify directed tissue movement along the anterior–posterior (AP) axis, we calculated the net AP displacement for each cell as the final x-position relative to its starting position and displayed the values in a boxplot.

### T1 Transition Analysis

To illustrate and quantify T1 transitions during germband extension, confocal timelapse movies of embryos were analyzed using TA [Aigouy et al., 2016]. Movies consisting of 100 timepoints, corresponding to 25 minutes of imaging from the onset of germband extension were selected. ROIs were defined on one side of the germband, excluding cells at the ventral midline. To minimize the effects of embryo movement, image registration was performed by fixing a tricellular junction near the cephalic furrow as a reference in Pyjamas. T1 transitions were initially detected automatically by TA and subsequently corrected and verified manually.

T1 transition frequencies were visualized using stacked bar plots showing the cumulative number of transitions occurring per minute for control and LARIAT conditions. Additionally, the average number of T1 transitions per minute was calculated for each movie and compared statistically between conditions using a linear mixed-effects model (LMM). The LMM included the experimental condition as a fixed effect and accounted for inter-movie variability by incorporating 'Embryo' as a random effect. Statistical significance was assessed by analyzing the fixed effect using an ANOVA F-test, and significance levels were indicated on plots.

### Apical constriction analysis

Cells and individual junctions were segmented in Pyjamas using time lapse movies and a watershed algorithm as described [Fernandez-Gonzalez and Zallen, 2011, Leung and Fernandez-Gonzalez, 2015, Fernandez-Gonzalez et al., 2021]. Automated segmentation was complemented by manual correction using the LiveWire tool (Pyjamas) if required. To compare the rates of apical area loss (Fig. 7A,B), mean apical areas were aggregated per experimental timepoint and embryo. Average areas were calculated across embryos at each time point, alongside SD and SEM, providing an estimate of variability across samples. We fitted an exponential decay model to quantify differences in the rate of area reduction between conditions and overlay the average area with. The decay model used was:

$$Area_{mean}(t) = A_0 \cdot e^{-kt}$$

where $A_0$ represents the initial mean apical area at the start of the fitted interval, $k$ is the decay rate constant, and $t$ is time in minutes. Model fitting was performed using nonlinear least squares regression (nlsLM in R).

Separate models were fitted to control and LARIAT conditions. Decay rate constants ($k$) were extracted and compared between conditions to quantify differences in apical constriction. The fitted exponential curves, along with SEM ribbons representing biological variability, were plotted to visualize and confirm the quality of fit and the precision of the estimated rates. To quantify decay rates per cell, an exponential decay model was fitted independently for each cell using the same exponential decay model as described above. Decay rates ($k$) were collected for each neuroblast and mesodermal cell and visualized using boxplots accompanied by jittered data points, highlighting distribution and variability across cells and experimental conditions. To compare fluorescence and area trajectories over time we used a Generalized Additive Model (GAM). Fluorescence values were normalized by subtracting image mode and dividing by image mean, then normalized again to initial values per cell. Time was scaled from 0 (start) to 1 (completion) to align data across cells. GAM smoothing was applied using cubic splines via the mgcv package in R, first individually per cell and then per experimental condition, highlighting overall trends.

**Anisotropic vertex model with dissipative adhesion**

We developed an anisotropic vertex-based model with dissipative junctional adhesion. We compare two experimental conditions: a control with normal E-cadherin expression and a LARIAT condition with altered E-cadherin levels. Differences in adhesion are encoded via distinct dissipative junctional adhesion coefficients (defined below).

The tissue is represented as a polygonal (cell) network where dynamics are governed by an anisotropic energy functional that accounts for the polarized tensions during germband extension [Wang et al., 2020, Erdemci-Tandogan et al., 2021] (Fig. S6A,B). The total energy is

$$E = \sum_{\alpha=1}^{N} [K_A(A_\alpha - A_{0\alpha})^2 + K_P(P_\alpha - P_{0\alpha})^2] + \sum_{\langle i,j \rangle} \gamma_{ij}\, l_{ij},$$

where the first sum runs over all cells and the second over edges connecting vertices $i$ and $j$. The area term penalizes deviations of the cell area $A_\alpha$ from its preferred value $A_{0\alpha}$. The perimeter term captures the competition between junctional adhesion and cortical contractility, penalizing deviations of the cell perimeter $P_\alpha$ from its preferred value $P_{0\alpha}$. $K_A$ and $K_P$ are the area and perimeter spring constants, respectively. To model polarized tension, the last term adds additional edge tension $\gamma_{ij}$ along edge length $l_{ij}$ between vertices i and j:

$$\gamma_{ij} = \gamma_0 \cos\left[2\left(\theta_{ij} - \varphi\right)\right],$$

where $\gamma_0$ sets the anisotropy amplitude, $\theta_{ij}$ is the edge angle, and $\varphi$ is the anisotropy direction which we set to $\varphi = \pi/2$.

**Dissipative junctional adhesion and substrate friction:** E-cadherin–mediated cell–cell adhesion is incorporated through dissipative terms in the equations of motion. Two friction sources are included: (i) substrate friction

$$\mathbf{f}_i^{\mathrm{S}} = -\mu\,\mathbf{v}_i,$$

and (ii) dissipative junctional adhesion

$$\mathbf{f}_i^{\mathrm{JD}} = -\zeta \sum_{j\in\mathcal{N}_i} \left(\mathbf{v}_i - \mathbf{v}_j\right),$$

where $\mathbf{v}_i$ and $\mathbf{v}_j$ are the velocities of vertices $i$ and $j$, $\mathcal{N}_i$ is the set of neighboring vertices connected to vertex $i$, $\mu$ is the substrate friction coefficient, and $\zeta$ is the dissipative junctional adhesion coefficient. This friction term is proportional to the relative velocity ($\mathbf{v}_i - \mathbf{v}_j$) between adjacent vertices, resisting their relative motion. In standard vertex models [Bi et al., 2015] $\zeta = 0$ and adhesion is incorporated only through the preferred perimeter $P_0$.

**Overdamped dynamics:** In the overdamped regime (negligible inertia), the net force on each vertex balances friction forces,

$$\mathbf{F}_i + \mathbf{f}_i^{\mathrm{fluct}} + \mathbf{f}_i^{\mathrm{S}} + \mathbf{f}_i^{\mathrm{JD}} = \mathbf{0},$$

where $\mathbf{F}_i = -\nabla_i \mathrm{E}$ is the force in vertex i and $\mathbf{f}_i^{\mathrm{fluct}}$ represents stochastic fluctuations. For a threefold coordinated network (each vertex has three neighbors),

$$\mathbf{F}_i + \mathbf{f}_i^{\mathrm{fluct}} - (\mu + 3\zeta)\,\mathbf{v}_i + \zeta \sum_{j\in\mathcal{N}_i} \mathbf{v}_j = \mathbf{0}.$$

We employ an Euler method to update the vertex positions,

$$\Delta \mathbf{r}_i = \mu_{\text{eff}} \mathbf{F}_i \Delta t + \boldsymbol{\eta}_i + \mu_{\text{eff}} \zeta \sum_{j \in \mathcal{N}_i} \Delta \mathbf{r}_j ,$$

where $\mu_{eff} = \frac{1}{\mu + 3\zeta}$ is the effective mobility and $\boldsymbol{\eta}_i$ is Gaussian noise with zero mean and variance $2\mu_{eff} T \Delta t$. The temperature $T$ introduces Brownian fluctuations [Brańka and Heyes, 1999].

**Simulation implementation:** Our implementation was built upon the open-source framework cellGPU [Sussman, 2017], extended to include box degrees of freedom (to capture tissue elongation) and dissipative adhesion terms. We initialized simulations with 256 cells in a periodic box. Initial cell configurations were generated using random Voronoi tessellations. The natural unit length of the simulations is given by $l = \sqrt{A_0}$ . We set the integration time step $\Delta t = 0.01\tau$, where $\tau$ is the natural time unit of the simulations. All simulations were run at their target parameters for $10^3\ \tau$ before activating the polarized tension (onset of tissue elongation).

**Model Parameters:** For the standard vertex model, we set the preferred shape index to $P_0 = 3.6$ (baseline) or $P_0 = 3.8$ (representing increased adhesion). For the dissipative vertex model, the preferred shape index was maintained at $P_0 = 3.6$ for both control and clustering conditions, based on experimental measurements showing no significant difference in cell shape index between control and Ecad-LARIAT embryos at the onset of germband extension (control: $3.99 \pm 0.02$, (n=4 embryos); LARIAT: $4.00 \pm 0.03$, (n=3 embryos); p=0.63, Mann-Whitney U test). Junction viscosity was set to $\zeta$=0.05 for control and $\zeta$=0.3 for clustering conditions, representing the increased frictional resistance in LARIAT embryos (Fig. S6C,D).

In both model frameworks, polarized tension was set to $\gamma_0$=0.3 for control and $\gamma_0 = 0.28$ for clustering, reflecting the minimal decrease in planar-polarized Myosin intensity (Fig. 4D). While reducing polarized tension alone affects tissue extension and T1 rates, this change is insufficient to reproduce the degree of tissue rigidification observed with E-cadherin clustering (Fig. S6E,F).

Common parameters across all simulations included preferred cell area $A_0 = 1$, area stiffness $K_A = 1$, perimeter stiffness $K_P = 1$, substrate friction $\mu = 1$, and temperature T = 0.02 in natural units.

n = 80 simulations per condition. Tissue extension rate was quantified from the time required for tissue length to double in the x-direction. T1 transitions were counted over the time window corresponding to control tissue doubling for comparison across all conditions.

**Measuring junctional tension:** We calculated junctional tension for any edge from:

$$T_{ab} = \frac{\partial E}{\partial l_{ab}} = 2K_P\left(\left(P_j - P_{0j}\right) + \left(P_k - P_{0k}\right)\right)$$

where $T_{ab}$ is the tension along the edge connecting vertices a and b at the interface between cells j and k [Yang et al., 2017].

**Statistical Analysis**

Statistical tests were chosen based on the distribution and structure of each dataset. For comparisons between two groups, parametric data were analyzed using Welch's t tests, whereas non-parametric data were analyzed using Wilcoxon rank-sum tests (same as Mann-Whitney U-test). For comparisons involving more than two groups, we used a Kruskal-Wallis test followed by Dunn's multiple-comparisons test with Bonferroni correction. For nested or repeated-measures datasets, in which multiple measurements were obtained from the same embryo or movie, we used linear mixed-effects models to account for non-independence by including condition as a fixed effect and embryo or movie as a random effect. Pairwise post hoc comparisons following linear mixed-effects models were performed using Tukey tests based on estimated marginal means. Statistical significance was defined as $p < 0.05$ and denoted as follows: * for $p < 0.05$, ** for $p < 0.01$, and *** for $p < 0.001$.

**Data processing in R**

Statistical analysis, modeling, and visualization were performed in R using packages from the tidyverse ecosystem, including dplyr, tidyr, ggplot2, purrr, and readr, together with ggpubr for statistical annotations and enhanced plotting. Additional visualization and figure-assembly tools included gridExtra, ggrepel, ggforce, plotrix, ggsci, scales, and circular. Nonlinear model fitting was carried out using minpack.lm and nls2, generalized additive models were implemented with mgcv, and linear mixed-effects models were fitted using lme4 and lmerTest, with broom.mixed used to tidy and summarize mixed-model outputs.

**Table S1**

| | **Source** | **Identifier** |
|---|---|---|
| Rat anti-E-cadherin | DHSB | DCAD2 |
| Mouse anti-Dlg | DHSB | 4F3 |
| Rat anti-Crb | Pellikka et al., 2002 | N/A |
| anti-GFP 488 (polyclonal) | Thermo Fisher | A-21311 |
| Mouse anti-Arm | DHSB | N27A1 |
| Guinea pig anti-αCat | Sarpal et al., 2012 | p121 |
| Mouse anti-Engrailed | DHSB | 4D9 |
| Phalloidin 647 | Thermo Fisher | A-22287 |
| Goat anti mouse Alexa 405 | Thermo Fisher | A-31553 |
| Goat anti guinea pig Alexa 555 | Thermo Fisher | A-21435 |
| Goat anti rat Alexa 647 | Thermo Fisher | A-21247 |
| D. melanogaster: “wild type” w1118 | BDSC | 3605 |
| D. melanogaster: w-RNAi | BDSC | 2640 |
| D. melanogaster: UAS-myr::GFP | N/A | N/A |
| D. melanogaster: matαtub67-Gal4 | D. St. Johnston | N/A |
| D. melanogaster: endo-Ecad::GFP | BDSC | 60584 |
| D. melanogaster: sqh-Sqh::mCherry | Martin et al., 2009 | N/A |
| D. melanogaster: Vinc::mCherry | Kale et al., 2018 | N/A |
| D. melanogaster: UAS-Ecad::GFP | BDSC | 58445 |
| D. melanogaster: UAS-Ecad RNAi | BDSC | 27689 |
| D. melanogaster: ubi-Ecad::GFP | Oda and Tsukita, 2001 | N/A |
| D. melanogaster: UASp-CRY2::vhhGFP | Gift from X. Wang | N/A |
| D. melanogaster: UASp-CIBN::MP | Gift from X. Wang | N/A |
| D. melanogaster: UASp-LARIAT | Gift from X. Wang | N/A |
| D. melanogaster: PH::PLCδ::GFP | Gervais et al., 2008 | 111 |
| D. melanogaster: αCat::YFP | KDSC | CPTI002516 |
| D. melanogaster: Scrib::GFP | BDSC | 94708 |
| D. melanogaster: sqh-Sqh::GFP | Royou et al., 2004 | N/A |
| D. melanogaster: aPKC::GFP | D. St. Johnston | N/A |
| D. melanogaster: PH::mCherry | Herszterg et al., 2013 | N/A |
| D. melanogaster: αCat::[MIMIC]K-RFP | S. Simoes, unpublished | N/A |
| FIJI | Schindelin et al., 2012 | RRID: SCR_002285 |
| Tissue Analyzer | Aigouy et al., 2016 | https://github.com/baigouy/tissue_analyzer |
| Pyjamas | Fernandez-Gonzalez et al., 2022 | https://bitbucket.org/rfg_lab/pyjamas/src/master/ |

### Supplementary Figure legends

**Figure S1: LARIAT-induced clustering of E-cadherin**

**(A)** Whole-embryo images (Fire LUT) comparing control and the Ecad-LARIAT embryos at 0, 15 and 30 minutes after onset of germband extension. Scale bar, 20µm.

**Figure S2: Ecad-LARIAT but not Ecad overexpression causes sustained high protein levels and morphogenetic defects.**

**(A)** Schematic of Drosophila germband extension. Stills and quantifications were taken within the first 30 minutes of germband extension.

**(B)** Illustration of the hypothesized difference between Ecad-LARIAT embryos and E-cadherin overexpression (Ecad-OE). Ecad-OE embryos compensate for high levels of E-cadherin by increased endocytosis, leading to no developmental effects, whereas in Ecad-LARIAT embryos stable E-cadherin clusters are arrested in the membrane and cause developmental defects.

**(C)** Still images of Ecad::GFP in control (endo-Ecad::GFP) embryos, Ecad-OE embryos (ubi-Ecad::GFP and UAS-Ecad::GFP), and Ecad-LARIAT embryos. Images were acquired from the same embryo per condition at 10-min intervals beginning at stage 7. Red arrowheads indicate cytoplasmic vesicles containing Ecad::GFP. Scale bar, 5µm.

**(D)** Quantification of normalized fluorescence of junctional and cytoplasmic E-cadherin levels in control, Ecad-OE, and Ecad-LARIAT embryos. Junctional E-cadherin progressively accumulates in Ecad-LARIAT embryos, whereas membrane signal in Ecad-OE embryos plateaus at much lower levels. Cytoplasmic E-cadherin is elevated in Ecad-OE embryos and declines as membrane levels decrease. $t = 0$ marks the onset of gastrulation. Sample sizes: control, 6 embryos, 41 cells; Ecad-OE, 6 embryos, 36 cells; Ecad-LARIAT, 7 embryos, 42 cells.

**(E)** Tissue flow during germband extension based on trajectories of tracked cells for 30 minutes. Individual cell tracks and the mean trajectory per condition are shown on the left. Anterior-posterior (AP) displacement was quantified for each tracked cell. AP displacement is significantly reduced in Ecad-LARIAT embryos compared with both control and Ecad-OE conditions, indicating impaired tissue movement. Sample sizes: control, 12 cells/embryos; Ecad-OE, 6 cells/embryos; Ecad-LARIAT, 11 cells/embryos. Statistics: Kruskal-Wallis test followed by pairwise Wilcoxon rank-sum tests with Bonferroni correction; control vs Ecad-OE, n.s.; control vs Ecad-LARIAT, $p = 0.0051$; Ecad-OE vs Ecad-LARIAT, $p = 0.0142$.

**Figure S3: E-cadherin cluster mobility is impaired in Ecad-LARIAT embryos.**

**(A)** Kymograph of individual E-cadherin clusters in the cell membrane over a 10-minute period, aligned at a tricellular (fixed) junction, shows low mobility in Ecad-LARIAT embryos. Scale bar, 1 μm. Cluster displacement was quantified as the maximum x-axis position over time.

**(B)** Average cluster displacement velocity and cumulative cluster displacement show that enhanced E-cadherin clustering impairs lateral mobility of E-cadherin clusters. Sample sizes: control, 6 embryos, 18 clusters; Ecad-LARIAT, 6 embryos, 18 clusters. Statistics: Wilcoxon rank-sum test; displacement velocity, $p = 0.00015$; cumulative displacement, $p = 0.00015$.

**Figure S4: E-cadherin co-localizes with interaction partners in Ecad-LARIAT embryos.**

**(A)** Zygotic LARIAT expression using *engrailed-GAL4* in endo-Ecad::GFP embryos shows enrichment of E-cadherin and Armadillo (Arm; = β-catenin) at cell contacts in Engrailed (En) domains. Scale bars: 20 μm.

**(B)** SIM imaging shows E-cadherin clusters at the En domain border, between Ecad-LARIAT expressing and non-expressing cells (right). Cell contacts between En positive and En negative cells show intermediate enrichment of Ecad::GFP. Scale bars: 2 μm.

**(C)** Colocalization between E-cadherin and junctional components in control and Ecad-LARIAT embryos at stage 8, ventral ectoderm. Scale bar: 5 μm.

**Figure S5: Myosin localization and dynamics in Ecad-LARIAT embryos.**

**(A)** Panels show the distribution of Ecad::GFP and myosin II (Sqh::mCherry) at multiple time points after onset of germband extension in control and Ecad-LARIAT embryos. Live-imaging snapshots were acquired every 5 minutes, Scale bar 5 μm.

**(B)** FRAP analysis of Sqh::mCherry (myosin II) shows no effect of Ecad-LARIAT on myosin II turnover. Half-time recovery or mobile fraction of myosin did not show significant differences between control and Ecad-LARIAT junctions (Half-time recovery, control: $1.28 \pm 0.214$ min, $n = 18$; LARIAT: $0.863 \pm 0.101$ min, $n = 17$; Wilcoxon rank-sum test, $p = 0.382$) (Mobile fraction, control: 0.651 ± 0.0540, n = 13; LARIAT: 0.728 ± 0.0444, n = 12; Wilcoxon rank-sum test, p = 0.430).

**(C)** Left: Z-projection of Sqh::mCherry (= myosin II) at stage 7, overlaid with vectors of medial myosin II displacement from cells outlined in blue and surrounding tissue. Vectors inside cells (not at junctions) were used for medial myosin flow quantification. Orientation: anterior left, dorsal up. Scale bar: 5μm. Middle: Instantaneous speed of medial myosin II in control and Ecad-LARIAT embryos (one frame per 5 s). Each value is the mean particle speed per frame. Control: 988 frames from 31 cells in 3 embryos; LARIAT: 1011 frames from 30 cells in 5 embryos. Median instantaneous speed (thick dashed lines) control/LARIAT: 0.045/0.050 μm $s^{-1}$; IQR (thin dashed lines) control/LARIAT: 0.038–0.059 / 0.036–0.063 μm $s^{-1}$. *** $p < 0.0001$ (Mann–Whitney test). Right: Mean instantaneous speed ratio along anterior-posterior (u, 0°) vs. dorsal-ventral axis (v, 90°). Mean control/LARIAT: 5.24/4.20. The dashed line marks no polarity (|u|/|v| = 1). A, anterior; P, posterior; D, dorsal; V, ventral. Mean ± SEM. * $p = 0.0231$ (Mann–Whitney test).

**Figure S6. Dissipative anisotropic vertex model framework related data and simulations.**

**(A)** Schematic illustration of the dissipative anisotropic vertex model showing tissue geometry and force components. Tissue represented as a polygonal cell network with vertices (grey circles) and cell-cell junctions (edges). A cell (blue) is highlighted showing cell area $A_\alpha$ and perimeter $P_\alpha$. Darker edges indicate junctions with higher polarized tension along the anterior-posterior axis. Black circle highlights vertex $i$ and its three neighbouring $j$ vertices. Vertex velocities $\mathbf{v_i}$ and $\mathbf{v_j}$ illustrate motion of the adjacent vertices.

**(B)** Mechanical forces acting on vertex $i$. For each vertex $i$, the net force $\mathbf{F}_i$ is derived from the mechanical energy $E$, which includes area elasticity, perimeter elasticity, and polarized tension terms (see Methods). Equation of motion for overdamped vertex dynamics shows the balance between energy-derived forces $\mathbf{F}_i$, substrate friction ($\mu\,\mathbf{v}_i$ ,), stochastic fluctuations ($\mathbf{f}_i^{\mathrm{fluct}}$), and dissipative junctional adhesion ($\mathbf{f}_i^{\mathrm{JD}}$) (see Methods). This friction term is proportional to the relative velocity ($\mathbf{v_i} - \mathbf{v_j}$) between adjacent vertices.

**(C,D)** Mean junctional tension in standard and dissipative vertex models. Mean edge tension averaged over all junctions and time for standard vertex model ($P_0 = 3.6$ vs 3.8)(C) and dissipative vertex model (ζ = 0.05 vs 0.3)(D) in natural simulation units. All other parameters are as defined in (Fig. 6). n = 80 simulations per condition. Box plots show mean (central line), box edges extend to ±1 SEM, and whiskers extend to ±1 SD, Mann-Whitney U test, *** $p < 0.001$.

**(E,F)** Polarized tension reduction alone is insufficient to explain changes in T1 transition frequency in Ecad-LARIAT conditions. Number of T1 transitions over time (E, left) and T1 transition rate per simulation time (right) for polarized tension $\gamma_0 = 0.3$ (control) and $\gamma_0 = 0.28$ (reduced tension). Bar plot shows mean and SEM. Tissue flow rate for both conditions. (F) Minimally reducing polarized tension alone modestly affects tissue extension and T1 rates but is insufficient to reproduce the degree of tissue rigidification observed with enhanced E-cadherin clustering (Fig. 5). Simulations performed with constant $P_0 = 3.6$ and $\zeta = 0$ (no junction friction). n = 80 per condition. Box plot shows mean (central line), box edges extend to ±1 SEM, and whiskers extend to ±1 SD. Time units in $\tau$. Other parameters: $A_0 = 1, K_A = 1, K_P = 1, \mu = 1$ and T=0.02 in natural simulation units. Mann-Whitney U test, T1 transitions (E) $p$=0.12, Tissue flow rate (F) $p$=0.11, both n.s.

**Video captions**

**Video S1:** 40x overview shots of Control (left) and Ecad-LARIAT (right) embryos expressing endogenously GFP-tagged E-cadherin. Video displayed at 14 frames per second.

**Video S2:** Cluster mobility at individual bicellular junctions in Control (left) and Ecad-LARIAT (right) embryos. Video displayed at 14 frames per second.

**Video S3:** Laser ablation in Control and Ecad-LARIAT embryos expressing Ecad::GFP. Timestamp, sec.ms, Images were taken every 2.3s with a 100x objective and movies are displayed at 14 fps.

**Video S4:** 5x movie of the first 300minutes of gastrulation captured with a brightfield microscope in Control (left) and Ecad-LARIAT (right) embryos. Frames were taken every minute. Video displayed at 14 frames per second.

**Video S5:** Germband extension (GBE) in control (left) and Ecad-LARIAT (right) embryos. Top panels show Ecad::GFP and bottom shows Sqh::mCherry labelling myosin. While myosin is not visibly affected, GBE is severely delayed upon clustering of E-cadherin. Video displayed at 14 frames per second.

**Video S6:** T1 transitions during germband extension in control and Ecad-LARIAT embryos. Each T1 transition was labelled in Tissue Analyzer (Aigouy et al., 2016).

**Video S7:** Standard anisotropic vertex model simulations. Simulation showing tissue dynamics for two preferred shape index values: $p_0 = 3.6$ (top, baseline) and $p_0 = 3.8$ (bottom, to account for increased adhesion). Higher shape index predicts enhanced T1 transitions and faster convergent extension, in contrast to experimental observations with enhanced E-cadherin clustering.

**Video S8:** Dissipative anisotropic vertex model simulations. Simulation showing tissue dynamics for two values of junction friction coefficient: $\zeta=0.05$ (top, control/wild-type) and $\zeta=0.3$ (bottom, clustering/LARIAT). Increasing junction viscosity substantially reduces T1 transition rates and slows convergent extension consistent with experimental measurements.

**Video S9:** Mesoderm apical constriction in Control (left) and Ecad-LARIAT (right) embryos. Top panels show Ecad::GFP and bottom shows Sqh::mCherry labelling myosin. Apical constriction dynamics are not perturbed in Ecad-LARIAT embryos. Video displayed at 14 frames per second.

**Video S10:** Neuroblast ingression in control (left) and Ecad-LARIAT (right) embryos. Apically constricting cells are outlined in green in an embryo expressing Ecad::GFP. LARIAT-clustering of E-cadherin delays apical constriction of neuroblasts. Embryos in both movies are between early stage 8 and early stage 9. Video displayed at 14 frames per second.

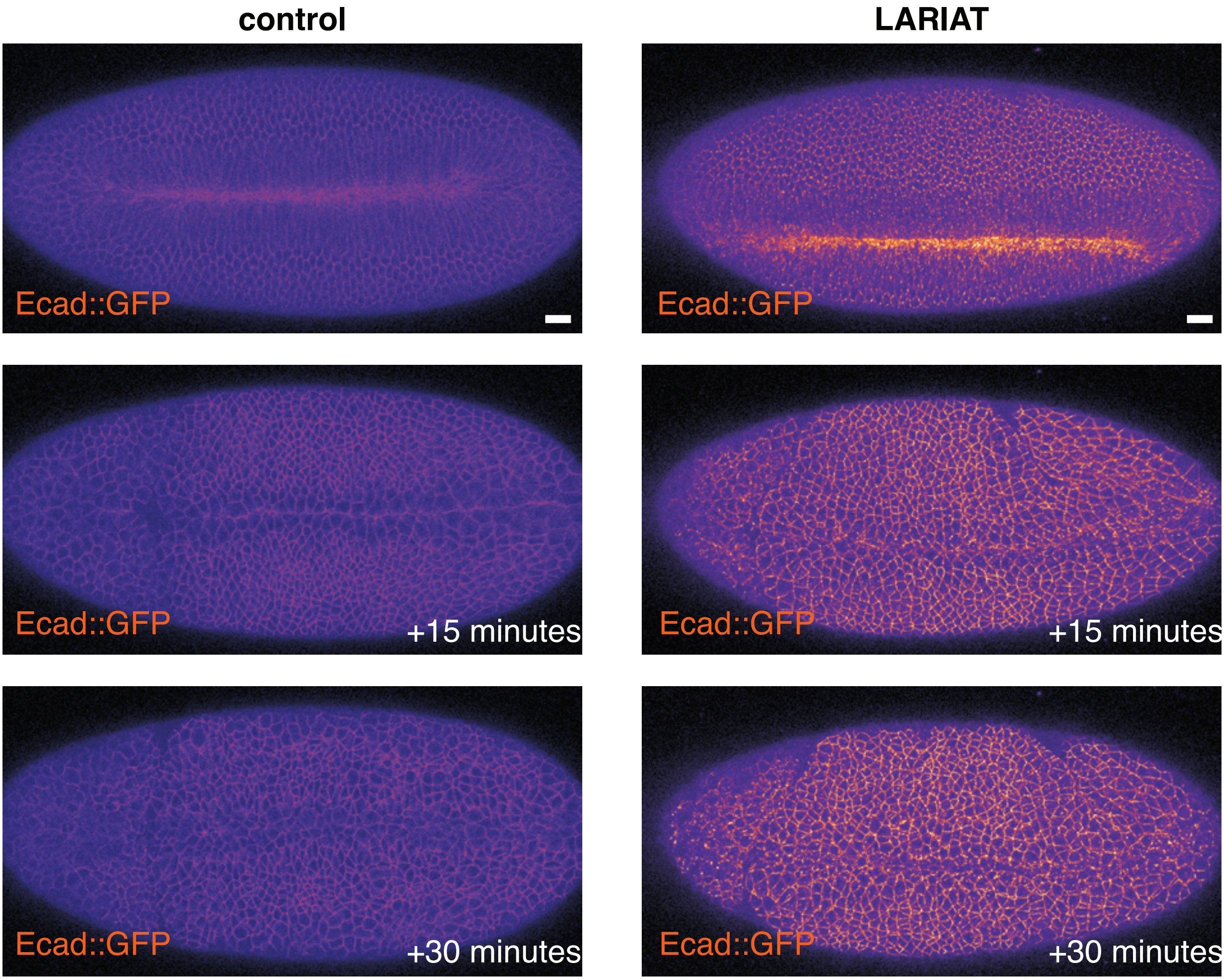


**Lerchbaumer et al., Figure S1**

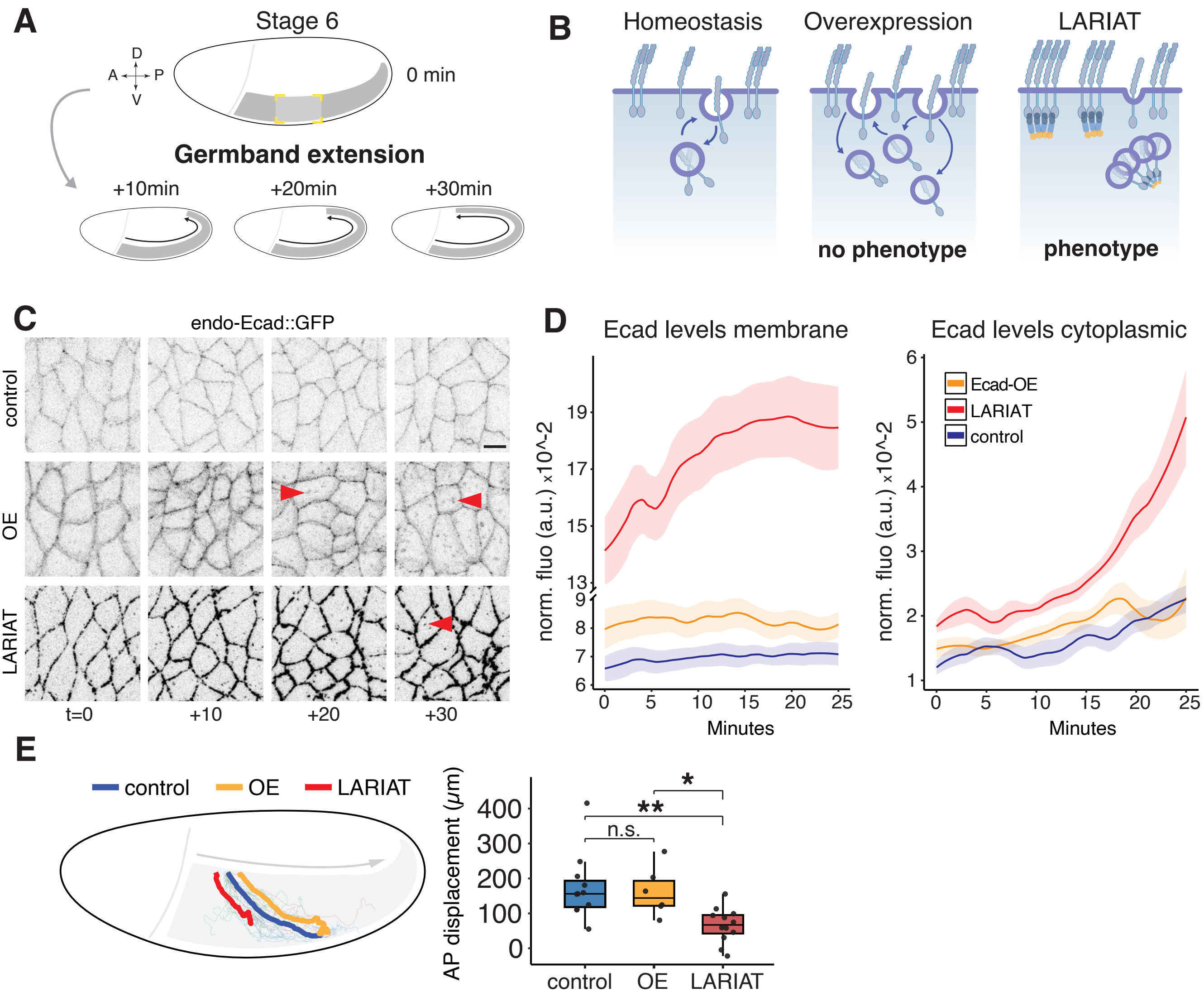


**Lerchbaumer et al., Figure S2**

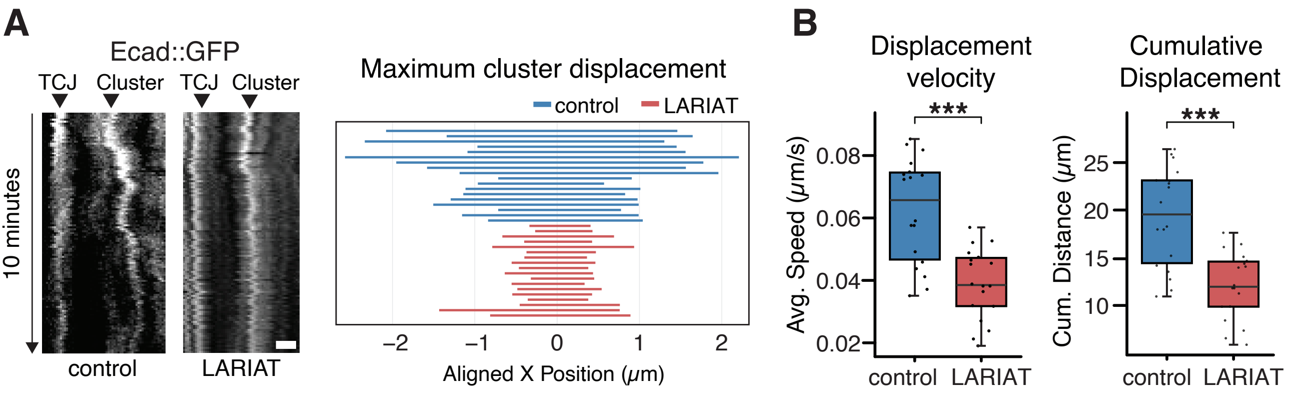


**Lerchbaumer et al., Figure S3**

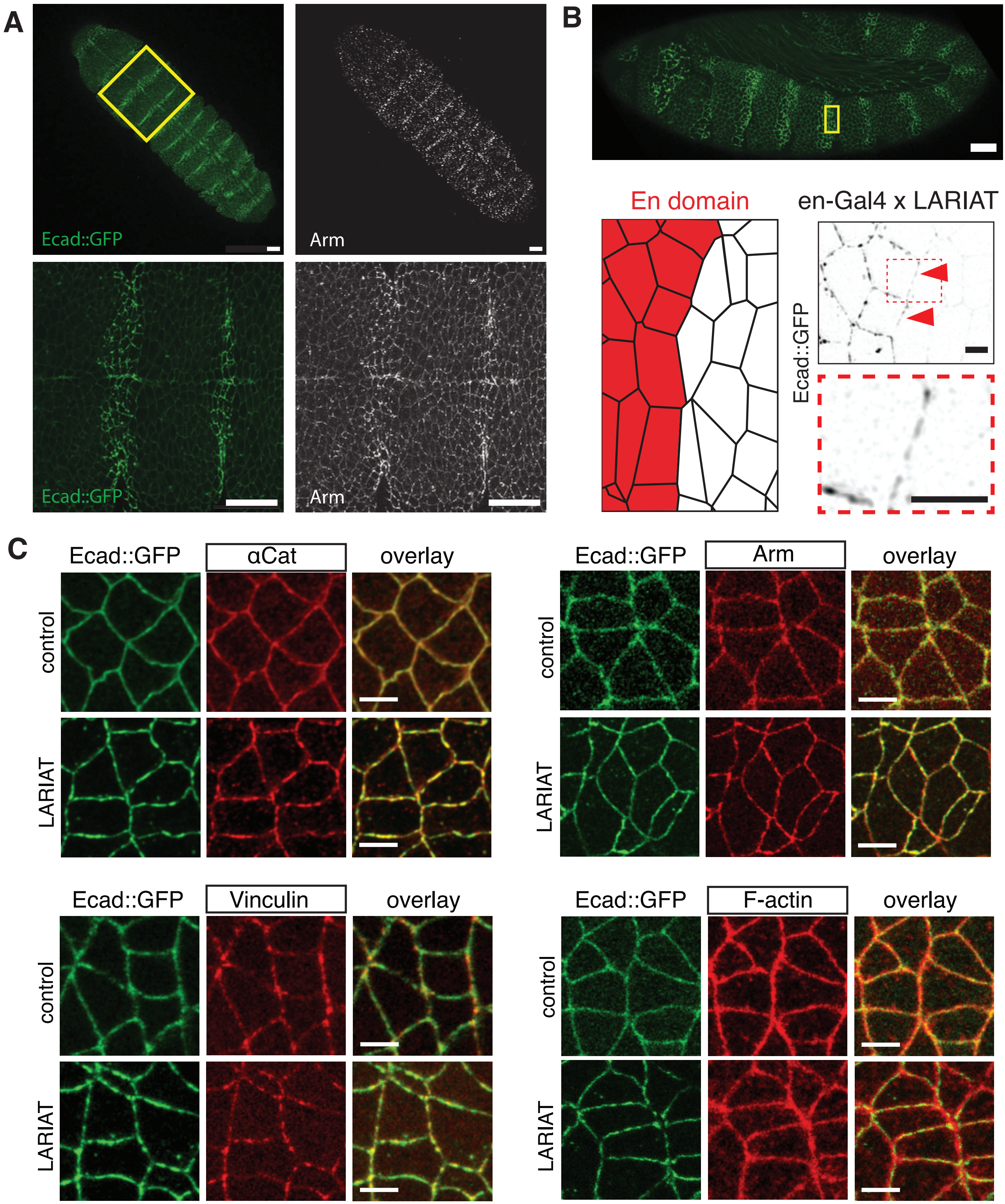


**Lerchbaumer et al., Figure S4**

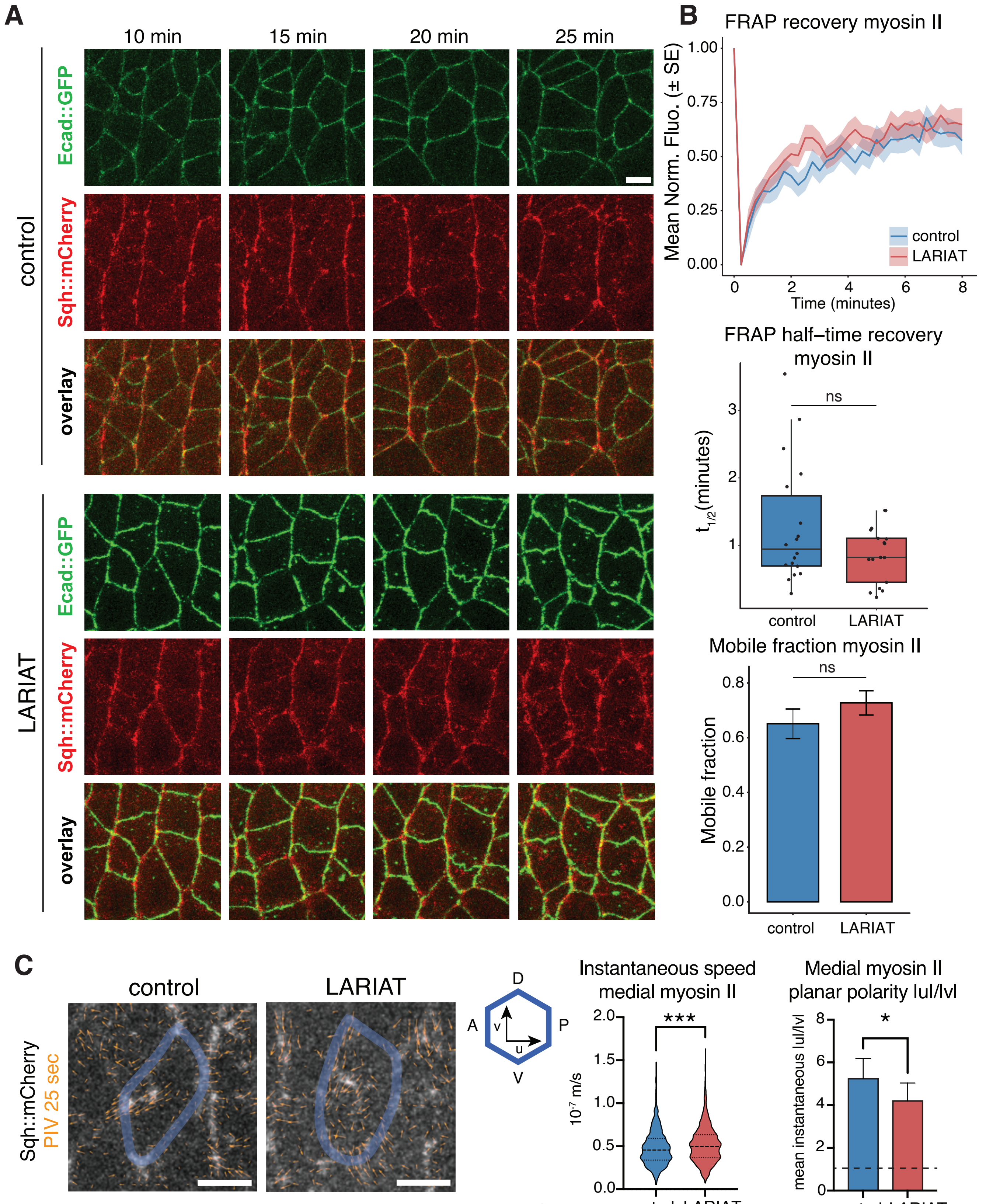


**Lerchbaumer et al., Figure S5**

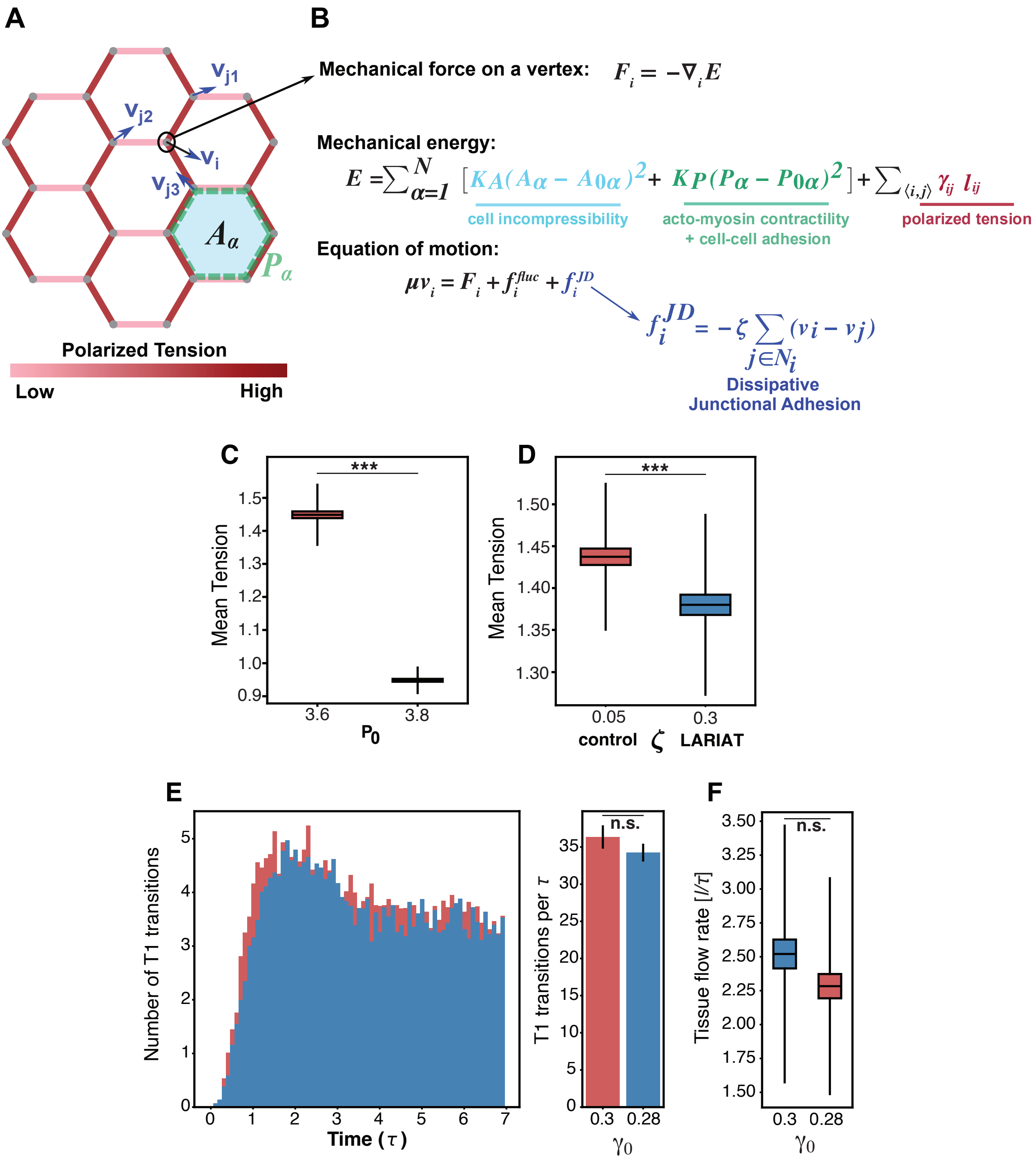


**Lerchbaumer et al., Figure S6**
